# Individual Differences in Cognition and Perception Predict Neural Processing of Speech in Noise for Audiometrically Normal Listeners

**DOI:** 10.1101/2024.08.28.609798

**Authors:** Sana Shehabi, Daniel C. Comstock, Kelsey Mankel, Brett M. Bormann, Soukhin Das, Hilary Brodie, Doron Sagiv, Lee M. Miller

## Abstract

Individuals with normal hearing exhibit considerable variability in their capacity to understand speech in noisy environments. Previous research suggests the cause of this variance may be due to individual differences in cognition and auditory perception. To investigate the impact of cognitive and perceptual differences on speech comprehension, 25 adult human participants with normal hearing completed numerous cognitive and psychoacoustic tasks including the Flanker, Stroop, Trail Making, Reading Span, and temporal fine structure (TFS) tests. They also completed a continuous multi-talker spatial attention task while neural activity was recorded using electroencephalography (EEG). The auditory cortical N1 response was extracted as a measure of neural speech encoding during continuous speech listening using an engineered “chirped-speech” (Cheech) stimulus. We compared N1 component morphologies of target and masker speech stimuli to assess neural correlates of attentional gains while listening to concurrently played short story narratives. Performance on cognitive and psychoacoustic tasks were used to predict N1 component amplitude differences between attended and unattended speech using multiple regression. Results show inhibitory control and working memory abilities can predict N1 amplitude differences between the target and masker stories. Interestingly, none of the cognitive and psychoacoustic predictors correlated with behavioral speech-in-noise listening performance in the attention task, suggesting that neural measures may be more sensitive measures of cognitive and auditory processing as compared to behavioral measures alone.

**SIGNIFICANCE STATEMENT:** These findings contribute to our understanding of how cognition affects the neural mechanisms of speech perception. Specifically, our results highlight the complex interplay between cognitive abilities and neural encoding of speech in challenging listening environments with multiple speakers. By incorporating these additional measures of cognition, we can achieve a more comprehensive understanding of an individual’s speech perception abilities, even in individuals with normal hearing. This approach could lead to earlier detection of hearing issues and more personalized interventions, ultimately enhancing communication outcomes for those with hearing difficulty.

## INTRODUCTION

Speech perception is a complex and multileveled process that requires the coordination of numerous neural systems, from decoding basic acoustic features of speech to transforming them into meaningful linguistic representations. Comprehension in noisy environments presents an additional challenge, placing substantial demands on every level of processing. This includes not only the encoding of stimuli and linguistic representations but also the segregation of auditory information into distinct streams (Griffiths & Warren, 2004), the ability to focus on a specific talker while filtering out others (Alain & Arnott, 2000; Shinn-Cunningham, 2008), and the capacity to maintain information over time (Martin, 2021). Consequently, performance in speech perception in noise is linked to both auditory and domain-general cognitive mechanisms (Akeroyd, 2008; Holder et al., 2018; Zekveld et al., 2013).

Speech perception in noisy environments disproportionately challenges listeners with hearing loss. However, many individuals with normal audiometric sensitivity also struggle with speech comprehension in such conditions. In fact, as many as 15% of individuals seeking hearing assistance at audiology clinics have normal audiometric thresholds but identify difficulty understanding speech in noisy settings as their primary concern (Cooper & Gates, 1991; Hind et al., 2011; Tremblay et al., 2015). Even among listeners with audiometrically normal thresholds, significant variability in speech-in-noise recognition can still be observed. For example, Ruggles & Shinn-Cunningham (2011) reported a range of correct response rates between 40-85% when participants were asked to report digits spoken by the target speaker in a simulated cocktail party scenario.

This variability in speech-in-noise (SiN) understanding could be influenced by individual differences in cognition and general perceptual ability. Previous work suggests the most critical cognitive mechanisms for SiN performance include selective attention (Shinn-Cunningham, 2008), working memory (Akeroyd, 2008; Gordon-Salant & Fitzgibbons, 1997; Ronnberg et al., 2008), inhibitory control (Janse, 2012, Sommers & Danielson, 1999), and more general aspects of executive function (Perrone-Bertolotti et al., 2017). Deficiencies in any of these domains can significantly impair speech perception (Pichora-Fuller et al., 1995). In addition to cognition, non-audiometric psychoacoustic abilities are also important, particularly sensitivity to temporal fine structure (TFS). TFS sensitivity is essential for spatial hearing, auditory object segregation and streaming, listening in the gaps (“glimpsing”), and pitch perception (Moore, 2008). As a result, TFS sensitivity is crucial for analyzing complex, ecologically valid auditory scenes, especially when speech is presented with background noise (Füllgrabe et al., 2015; Oberfeld & Klöckner-Nowotny, 2016). Notably, Oberfeld & Klöckner-Nowotny (2016) found that both cognitive measures, including selective attention, and psychoacoustic measures, such as TFS sensitivity, can predict SiN performance in individuals with normal hearing. Thus, explorations of speech-in-noise recognition abilities must also account for individual differences in both cognitive and basic auditory skills.

Additionally, successful speech (in noise) recognition relies on robust neural encoding mechanisms within the auditory system. The N1 component, for example, reflects early cortical processing of sounds (Winkler et al., 2013). The N1 is considered a general index of early perceptual encoding of speech sounds, and it is sensitive to acoustic cues within a stimulus, speech or otherwise (Getz & Toscano, 2021). Additionally, N1 responses have been associated with auditory selective attention and stream segregation which are essential for distinguishing between different sound sources in the environment (Gutschalk et al., 2007; Snyder et al., 2006). Presumably, individual differences in speech perception should also be linked with variability of neural processes within the auditory cortex. Yet, despite the established influence of cognitive and non-audiometric psychoacoustic abilities on speech perception, how these individual differences impact the neural encoding of speech (and SiN; Bednar & Lalor, 2020) remains unclear. This knowledge gap limits our ability to fully explain the variability in speech perception abilities among normal-hearing individuals in noisy environments.

The goal of this study is to understand how cognitive factors known to be important for speech perception influence speech processing in the auditory cortex. We specifically focus on the differences in N1 amplitude between target (to-be-attended) and masker speech. By measuring N1 amplitudes in response to these conditions, we aim to understand the brain’s mechanisms for filtering and prioritizing relevant auditory information over irrelevant stimuli. We hypothesize that neural mechanisms involved in cocktail party listening are shaped not only by basic auditory factors, such as hearing sensitivity, but also by an individual’s cognitive abilities. Using multiple regression analysis, this study identifies which cognitive/psychoacoustic functions among individuals with normal audiometric hearing impact the neural mechanisms (specifically the N1 component) involved in attending to target speech and inhibiting masker speech. Collectively, these findings shed light on the cognitive contributions to neural mechanisms relevant for SiN perception.

## METHODS

### Participant Information

Twenty-nine participants were recruited for this study. Eligible participants were required to be between 18 and 40 years old, speak English as their first language, and report no neurological, psychiatric, or scalp conditions that could directly impair their ability to understand or attend to speech or hinder electrophysiological measurements (e.g., epilepsy, certain strokes, ADHD, scalp issues such as wounds). Participants were screened for visual acuity (< 20/40 per Snellen chart), normal hearing (<25 dB HL air conduction thresholds between 250-8000 Hz in both ears, no air-bone gaps >10 dB, no interaural asymmetries >20 dB at 500, 1000, or 2000 Hz), and cognitive abilities (>24 MoCA score) (Nasreddine et al., 2005).

One participant withdrew from the study due to personal reasons, two were excluded for failing the hearing screening at the time of testing, and one was excluded due to technical issues with data acquisition. Additionally, one participant was identified as an influential outlier in the Trail Making Test based on a Cook’s Distance test threshold of 0.5. This resulted in a final cohort of 24 participants. The average age of participants was 23.4 ± 5.02 SD years (range: 18-38 years). Fourteen participants were female and ten were male. Twenty-one participants self-identified as right-handed, one as left-handed, and two as ambidextrous. On average, participants had 17.00 ± 3.18 SD years of education and 4.21 ± 3.82 SD years of formal music training. Pure tone averages (PTA) were obtained for each ear as the average threshold at 0.5, 1, and 2 kHz. Average air conduction PTA thresholds were 6.88 ± 3.78 SD and 7.04 ± 4.56 SD for the right and left ears, respectively. All participants provided informed consent as approved by the university’s Institutional Review Board and were financially compensated for their participation and travel.

### Project Overview

Each participant completed three sessions in total: a 1-hour audiological exam, a 1.5-hour behavioral session, and a 2.5-hour laboratory EEG session. Most participants completed more than one session on the same day (e.g., audiological + behavioral or behavioral + EEG), with a long break in between to minimize fatigue. The analysis presented in this paper is based on a subset of data from a larger study investigating hearing across the lifespan, which will be detailed in future reports. Consequently, only a portion of the measurements and tests conducted during the study are included in this analysis.

### Audiology Session

Participants completed a standard audiologic assessment either conducted by a professional audiologist or by highly trained personnel in the lab. Before inserting earphones, an otoscopic examination was conducted to check for tympanic membrane pathologies or any obstructions in the ear canal (i.e., cerumen) that could prohibit the use of insert earphones. Participants found to have ear canal occlusions or other visible pathologies were advised to consult their primary care provider and return to the study once the issue was resolved.

Pure-tone audiometry was performed for both ears either at a local affiliated audiology clinic or in a quiet, sound-dampened room within the lab. Air-conduction thresholds were measured using pulsed pure tones at frequencies of 0.25, 0.5, 1, 2, 3, 4, 6, and 8 kHz with insert earphones. Bone-conduction thresholds were measured using a RadioEar bone conductor placed on the mastoid at 0.25, 0.5, 1, 2, 3, and 4 kHz. The thresholds were determined using a 10 dB down/5 dB up staircase procedure. Pure tone averages (PTA) were obtained for each ear as the average air-conduction threshold at 0.5, 1, and 2 kHz. While several other tasks typical of a standard audiology exam were conducted, only air and bone conduction measures are described here as they were the indicators used to determine normal hearing in participants.

### Cognitive Tasks

Cognitive tasks were administered in a randomized order for each participant.

#### 1. Flanker Test

The Flanker test measures visuospatial interference, selective attention, and inhibitory control. We used a computerized version of the task using the PEBL software (Psychology Experiment Building Language; Mueller & Piper, 2014). In the task, five light gray arrows were displayed in the center of the screen against a black background. The arrows were either all pointing in the same direction (congruent condition) or with the center arrow pointing in the opposite direction to the flanking arrows (incongruent condition). Participants were asked to press the left or right shift keys on a keyboard corresponding to the direction of the center arrow. A fixation cross appeared for 500 ms before the arrows were presented, and participants had 2 seconds to respond before the trial timed out. A blank screen was shown for 2 seconds before the next trial began. Participants completed 8 practice trials, with two trials for each combination of conditions (left/right, incongruent/congruent), followed by 80 test trials (20 per condition). The interference effect was calculated as the median reaction time difference between incongruent and congruent trials.

#### 2. Stroop

The Stroop test is a long-standing, well-established assessment of linguistic interference and response inhibition (Strauss et al., 2006; Stroop, 1935). In our study, we utilized a computerized version of the Victoria Stroop Test (VST; Strauss et al., 2006), implemented using PEBL software (Mueller & Piper, 2014). The VST was selected for our study due to its shorter test administration time compared to other standardized Stroop tests. Evidence suggests that shorter test durations may be more sensitive to individual differences on this task (Klein et al., 1997). The VST has been proven effective with a wide range of adults, from 18 to 94 years old (Troyer et al., 2006).

As with other Stroop versions, the VST involves color identification of items presented to the participant, including a color word interference task. The VST is divided into three blocks always presented in the following order: colored dots (Part D), neutral words (Part W), and color words presented in contrasting text colors (Part C). Each block displays 24 items in a 6 x 4 grid on a gray background. Items are colored red, blue, green, or yellow, and participants press a corresponding number key to identify the color of each item. A square is shown around the current item trial, and it only moves to the next item with a correct response. The square flashes white when an incorrect response is made. Each color is used six times, and the item colors and the color-number keypad pairs are pseudorandomized for every participant (one of each color in each row). Part D serves as a practice block for color-number mapping and is not scored here. For Part W (neutral words) and Part C (color words), participants must name the colors in which the words are printed, disregarding the verbal content printed in lowercase letters (neutral words: when, hard, and, over; color words: red, green, blue, yellow). In Part C, the color name never corresponds to the text color, thus requiring individuals to inhibit an automatic reading response and only identify the text color itself. The interference effect is measured as the difference in median reaction times between Part C (i.e., the linguistic interference task) and Part W (i.e., the control task).

#### 3. Reading span

Working memory ability was assessed using the reading span test (Daneman & Carpenter, 1980), as employed in previous research (Dryden et al., 2017; Lunner, 2003; Rudner et al., 2012). We administered a computerized version of the reading span via the Psychology Experiment Building Language (PEBL) program (Mueller & Piper, 2014). The PEBL reading span test was developed for automated scoring as described by Unsworth et al. (2009). In this task, participants read sentences displayed on the computer screen and indicated whether each sentence made sense by pressing a key (i.e., in grammatical, coherent, or plausibility sense). Sentence lengths ranged between 9 and 14 words. After the sentence, a letter was presented on the screen for 1 second, and participants were required to memorize the sequence of letters presented across sentence-letter pairs. The task consisted of blocks of 3 to 7 sentence-letter pairs, with each block length presented three times in random order throughout the test. At the end of each block, participants selected the correct sequence of letters from a 4 x 3 grid displayed on the screen and were instructed they could guess or leave a blank in the letter sequence if they were uncertain. They were then provided feedback regarding both sentence judgment and letter sequence accuracy.

Prior to the test, participants completed two practice examples each for sentence reading, letter memorization, and sentence-letter memorization pairs. The average reading time during these practice trials determined the timeout duration for the test blocks (i.e., timeout = 2.5 × average practice reading time) (Unsworth et al., 2009). If participants did not respond to a sentence before the timeout period, the sentence trial was marked incorrect, and the subsequent letter was shown on the screen. Throughout the test, sentences, letters, and feedback were shown in white text on a black screen background.

Working memory span was determined by calculating the average accuracy of letter recall for 3 and 4 letter trials, when any letters were correctly recalled in the correct order (i.e., each correct letter contributing to the score). Though earlier research has suggested a working memory capacity of up to ∼7 items (Miller, 1956), recent studies indicate that this capacity is lower, and can vary based on task demands and circumstances (Cowan, 2015). Consistent with findings indicating working memory capacity is approximately 4 items (for reviews, see Cowan, 2001, 2015), our findings revealed that the accuracy for the 3 and 4 letter trials provided more stable and informative data for our model compared to the average accuracy across 3 to 7 letter trials (pilot data not shown). Therefore, we used the average accuracy from the 3 and 4 letter trials as our primary measure of working memory span.

#### 4. Trail making test

The Trail Making Test (TMT) (Reitan, 1958) was used to assess cognitive flexibility and executive function. In TMT-A, participants connect lines sequentially between numbers, while TMT-B requires them to alternate between numbers and letters (e.g., 1-A-2-B). Participants were given TMT-A followed by TMT-B, with a short practice at the beginning of each. The test was administered via PEBL (Mueller & Piper, 2014), with white nodes displayed on a gray background and black text for letters and numbers. Participants clicked on the nodes in the correct sequence as quickly and accurately as possible, and lines were drawn on the screen to connect consecutive nodes upon correct selection. Given known limitations of the original Reitan-developed TMT A/B (i.e., length differences between the nodes of the A and B versions) (Gaudino et al., 1995), our version includes fixed node positions as mirrored images across TMT-A and B to ensure similar distance lengths for the two tests (Strelcyk et al., 2019). Any differences in reaction time are therefore presumed to reflect processing differences across the A and B versions. Executive function and cognitive flexibility were measured as the difference in median reaction times of TMT-B and TMT-A conditions.

#### 5. Temporal fine structure

The ability to localize low-frequency sounds along the horizontal plane relies on interaural time differences (ITD), which are important cues for distinguishing multiple sounds such as recognizing speech from a target talker amidst spatially separated background noise or other interfering sounds (Moore, 2021). For sinusoidal signals, ITDs are equivalent to interaural phase differences (IPDs), and the ability to discriminate changes in IPDs depends on binaural sensitivity to temporal fine structure (TFS). To assess IPD discrimination abilities, we used the TFS-LF (low-frequency) test (Hopkins & Moore, 2010), which is implemented in a computerized software program developed by Sęk and Moore (Sęk & Moore, 2012, 2021). In the TFS-LF test, participants are tasked with discriminating sinusoidal tone bursts with adaptively varying IPDs, where the tone envelopes are synchronized across both ears, ensuring that task performance relies on sensitivity to TFS (IPD). TFS-LF administration was consistent with previous work and is briefly described below (Füllgrabe & Moore, 2018; Hopkins & Moore, 2010; Sęk & Moore, 2012, 2021).

IPD discrimination was evaluated at test frequencies of 250 and 500 Hz, consistent with previous studies (for review, see Füllgrabe & Moore, 2018). Intensity level was set individually for each ear 30 dB above the participant’s pure tone thresholds at the tested frequencies (i.e., 30 dB sensation level [SL] at 250 and 500 Hz). The TFS-LF task consisted of a two-interval, two-alternative-forced-choice design, where each interval included four successive tones. In one of the intervals, all four tones had a consistent 0° IPD, while in the other interval, the IPD alternated between 0° and φ across the four successive tones. A 0° IPD is perceived as a tone localized to the center of the head, whereas a large IPD is perceived as a tone lateralized (or as more diffuse) toward one ear. Participants were asked to identify the interval in which the tones sounded different or appeared to change in some way. Each tone had a duration of 400 ms, with 20 ms rise and decay times and a 300 ms inter-interval gap between tone blocks. Feedback was provided after each trial. The IPD φ was adaptively varied, starting from 180° (maximally lateralized), using a 2-down, 1-up procedure to estimate a threshold corresponding to 71% correct responses. That is, the value of φ is divided by a factor k after two successive correct responses or multiplied by factor k following one incorrect response. Before the first turn point, k = 1.253; between the first and second turn points, k = 1.252; and k = 1.25 after the second turn point. The threshold was computed as the geometric mean value of φ at the last six turn points, and the final TFS-LF measure was then calculated as the average across both 250 and 500 Hz frequencies.

#### Additional tests

During the cognitive session, participants also completed two additional tests: 1) a pitch discrimination task and 2) a Speech Recognition in Noise Test (SPRINT), which assesses SiN perception. These measures were not included as predictors in the current analysis, as our primary focus was on evaluating variables that most closely align with existing research on predictors of SiN perception (Akeroyd, 2008; Janse, 2012; Perrone-Bertolotti et al., 2017; Shinn-Cunningham, 2008).

### Continuous Multi-Talker Spatial Attention Task

#### Stimuli

The stimuli consisted of short fairytale stories available in the public domain (available for listening on www.librivox.com). Stories were selected to have content that was unlikely to be widely recognized, reducing the chances of participants relying on modern renditions for comprehension recall (cf. stories popularized by children’s books or movies). Additionally, each story was chosen to have a similar narrative arc—i.e., up through rising action, climax, or early falling action— within a 7.5-minute block to ensure sufficient and consistent content for probing comprehension. Thirteen stories were piloted with four raters who evaluated them based on factors such as interest and engagement, ease of understanding, complexity of grammar or writing style, familiarity (i.e., low scores preferred), and suitability for an adult audience (not only for children). The seven highest-scoring stories across these criteria were selected for use in the experiment.

The short stories were revised to incorporate monosyllabic color words as targets for the experiment (e.g., red, green, black, white, etc.). The color words were integrated into the story itself rather than being inserted as unrelated, random target words (e.g., “…with a narrow *blue* ribbon over her shoulders”). A male native English speaker with an American Midwestern accent recorded the modified stories using a Shure KSM244 vocal microphone (cardioid polar pattern, high-pass filter at 18 dB per octave with an 80 Hz cutoff) and Adobe Audition (sample rate = 48,000 Hz, 32-bit depth [float]) in a soundproof, quiet room. Silent gaps in the recordings longer than 500 ms were shortened to 500 ms, following procedures similar to those in previous studies (Broderick et al., 2018; Teoh et al., 2022; Teoh & Lalor, 2019). Finally, the edited recordings were cropped to 7.5 minutes per story.

#### Cheech

The short story auditory stimuli were subsequently modified using a patented technique known as “Cheech” (“chirped-speech”), which is intended to elicit robust auditory evoked potentials (Backer et al., 2019; Miller et al., 2020). In this process, some of the glottal pulse energy within specified frequency bands is replaced with narrowband synthetic chirps (for more details, see Backer et al., 2019). These chirps are aligned with the natural glottal pulse energy in voiced speech, creating an acoustically fused auditory perceptual object. This modification maintains the naturalistic qualities and linguistic content of speech while introducing sufficient transient chirp activation to measure auditory brainstem, thalamic, and cortical responses.

The process begins by identifying voiced epochs where chirps will be inserted. Audio was filtered from 20 to 1000 Hz, and voiced periods of at least 50 ms were defined when the speech envelope between 20-40 Hz exceeded a threshold of approximately 28% of the overall speech root-mean-square (RMS) amplitude. The timing of the glottal pulses within these voiced periods was determined through a speech resynthesis process using custom MATLAB code (The MathWorks Inc., 2021) and the TANDEM-STRAIGHT toolbox, which retains the original speech periodicity characteristics (Kawahara et al., 1999, 2008; Kawahara & Morise, 2011). The continuous speech was then re-filtered into alternating, octave-wide frequency bands: 0-250, 500-1000, 2000-4000, and 11,000-∞ Hz, with chirp energy constrained to frequency bands of 250-500, 1000-2000, and 4000-11,000 Hz. The chirp and speech bands in the alternating, interleaved octave-wide bands were then added together. Each mono track was then duplicated to form stereo audio. Before the Cheech synthesis, audio files were normalized to ensure consistent chirp amplitudes relative to the speech levels across each story, and the RMS amplitudes were re-checked across all Cheech-modified stories to ensure equivalence after the Cheech process was completed.

The timing of the chirps varied with the natural fluctuations in the running speech, with special modifications to optimize measurement of auditory event-related potentials (AERPs). Specifically, when voicing energy was detected, chirps were inserted at minimum 18.2 ms apart (55 Hz), skipping glottal pulses that occur within the 18.2 ms window. The first chirp in any sequence was always followed by a minimum interval of 48 ms before the next chirp was presented, skipping glottal pulses in between. This longer ISI was designed to enhance the measurement of middle-latency responses (MLRs); we use this “first chirp” to elicit LLR N1s as well.

In the dual-talker conditions, one story from each pair was pitch and vocal-tract modulated using MATLAB STRAIGHT prior to the Cheech process. Specifically, stories intended to represent a female voice were pitch modulated to an average fundamental frequency (F0) of 180 Hz, while the original male voice audio had an average F0 of approximately 128 Hz. By modulating the F0, key speaker characteristics such as phrasing, intonation, and pacing were maintained, as all stories were initially recorded by a single talker. This pitch modification also introduced a combined voice-gender and spatial release from masking effect, which improved pilot test performance (Oh et al., 2021, 2022).

#### Continuous multi-talker spatial attention task

The Cheech stimuli were spatially filtered using head-related transfer functions (HRTFs) from the SADIE II database (Armstrong et al., 2018) to simulate audio at ±15°. This spatialized audio was mirrored by two symbols on two computer screens centered approximately 15° to the left and right of the participant’s midline. The symbols “<” and “>” indicated the target story’s location on the left or right, respectively, while a “+” symbol denoted the location of the target story, as shown in Figure 1. The visual symbols were identical in size, color, and line lengths to ensure consistent luminance for eye-tracking measurements (not analyzed in this report). Custom MATLAB Psychtoolbox code (Brainard, 1997; Pelli, 1997) was used to program the spatial and visual stimuli to switch between left and right locations 75 times over a 7.5-minute period, with each switch occurring approximately every six seconds (random distribution of 6 ± 1 seconds). The visual icons switched instantaneously at each switch cue, while the audio crossover fade time was set to 35 ms, starting at the onset of each switch cue. During piloting, this fade duration was found to effectively minimize audio artifacts, such as “pops,” while also reducing the perception of a gradual audio glide between spatial locations (i.e., as sudden of a jump as possible without introducing noticeable distortions).

**Figure 1:**
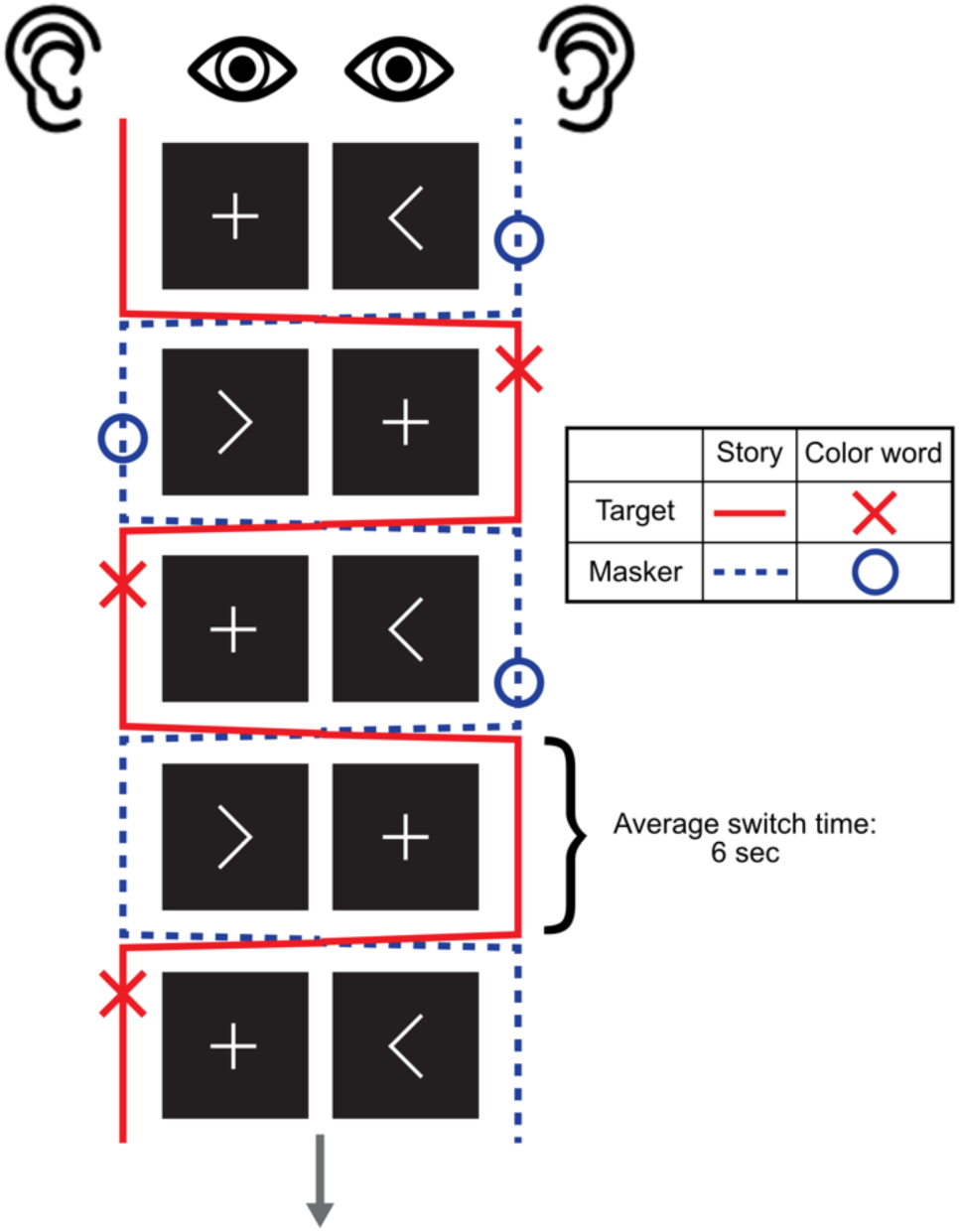
Continuous multi-talker spatial attention task procedure. The figure illustrates the visual cues presented on screen to participants during the task while they listened to a series of short stories. The target (and masker when present) alternate between spatial locations. In the mono-talker condition, only the target story is present, with its spatial location still switching sides. Participants are instructed to focus on the side of the screen indicated by the “+” symbol, which marks the location of the target story. They are also instructed to press the spacebar when they hear a color word spoken by the target speaker (red “X” symbols) and ignore any color words in the masker story (blue “O” symbols).

The target color words were randomly assigned to one of five possible onset intervals ranging from 0.125 and 2 seconds after a spatial switch cue (i.e., 0.125, 0.25, 0.5, 1, and 2 seconds; 5 color words per interval). These intervals were determined based on preliminary piloting that aimed to evaluate whether brief lapses in attention following a shift in spatial focus— analogous to "attentional blinks"— affected participants’ ability to detect color words; these results will be presented in a subsequent paper. Onset timings for the color words were calculated using Gentle, an audio forced aligner (https://lowerquality.com/gentle/). The closest preceding switch time was adjusted to match the assigned switch-to-color-word interval, resulting in 50 random switches and 25 switch-to-color-word events (i.e., a 33% probability of a color word occurring within 2 seconds of a switch). These 25 color words were evenly distributed between left-to-right and right-to-left switches within each story, minus the one remainder.

#### Experimental conditions

After cropping the audio, the seven stories were assigned to specific experimental conditions: two stories presented alone (“mono-talker”), two sets of paired stories (“dual-talker”), and one for initial practice. Half of the stories (one mono-talker and one dual-talker pair) had modified Cheech characteristics to study their effects on brainstem encoding (results forthcoming in future reports); however, this modification did not impact cortical responses, so we used all six non-practice stories for our N1 analyses. Since our focus is on predicting SiN performance, this paper primarily examines the dual-talker conditions (excluding Figure 4 and associated analyses). In the dual-talker paradigm, each story was presented twice—once as the "target story" and once as the "masker story" within its pair. This experimental manipulation allowed us to assess the effects of directed spatial attention, as participants were instructed to either "attend" to or "ignore" the same audio depending on its role in the trial. The order of stories and conditions was counterbalanced across participants using a Latin square design to minimize potential sequence-dependent learning effects. Additionally, no pair of stories was presented consecutively.

Story pairs in the dual-talker condition required additional considerations. As described in the Cheech section above, one voice in each story pair was pitch and vocal tract modulated to simulate a female speaker. Consistent with findings on the combined effects of voice-gender and spatial release from masking (Oh et al., 2021, 2022), two distinct talker F0s facilitated better listener differentiation between spatially separated voices and improved task performance during piloting. The mono-talker stories were presented in their original, unmodified male voice. To minimize the influence of stimulus variability on the N1, the analysis in this paper focused solely on the male target and masker conditions. Due to the naturalistic embedding of color words within the stories, some overlap occurred in the dual-talker pairings, i.e., color words in the target and masker speech occurred in close temporal proximity. Story pairs were selected to minimize color word overlaps, resulting in six overlapping words that were excluded from the behavioral analysis (four from one story pair and two from the other pair). A few target story switch times were adjusted to ensure that no color words in the masker story occurred during the switch (i.e., masker color word onset was not between -0.5 to 0.1 ms from a switch time).

#### Stimulus presentation equipment

Participants were seated in a soundproof, electromagnetically shielded booth approximately 176 cm from two computer monitors. The Dell UltraSharp monitors had a 60 Hz refresh rate with screen settings maintained at standard factory defaults. Stimulus presentation, including both audio and visual components, was controlled using custom MATLAB and Psychtoolbox code. Auditory stimuli were routed from a Hammerfall audio card to an RME Fireface UFX II audio interface, then through an RME ADI-2DAC FS headphone amplifier, and finally delivered via ER-2 insert earphones (Etymotic Research/Interacoustics). The ER-2 insert wires were electromagnetically shielded up to the transducer box to minimize EEG artifacts (Campbell et al., 2012). The Cheech stimuli were presented at 75 dB SPL.

In addition to the 2-channel audio tracks, a pair of channel outputs carrying trigger events was routed from the Fireface audio interface to two Brain Products StimTraks. These StimTraks monitored triggers for the target and masker stories, respectively, converting the audio trigger signals into pulses sent to a TriggerBox (Brain Products GmbH). This configuration ensured that audio playback was synchronized with the chirp triggers embedded during the Cheech process, minimizing latency jitter, and maintaining sample-level synchronicity. Triggers managed by Psychtoolbox—such as participant responses, block sequence triggers, and spatial location switches—were also routed to the TriggerBox through the computer parallel port. The TriggerBox then transmitted both parallel port and StimTrak triggers to the Biosemi signal receiver box (refer to EEG recordings).

#### Attention task procedure

The order of events within each block is as follows: a 1-minute rest period (for resting state EEG analysis), 7.5 minutes of audio presentation, comprehension questions, subjective workload questions, a flash reaction time task, and a break period. During the rest period, participants were instructed to remain still with their eyes open. They were instructed to attend to the target story as the auditory stream alternates between left and right spatial locations. They must immediately shift both their selective attention and eye gaze to the fixation point (“+”) on the computer screen as it switches the cue. Participants were also instructed to press the spacebar upon hearing the embedded color words in the story as quickly and accurately as possible. In dual-talker condition blocks, they were instructed to ignore color words from the masker story and only respond to those from the target story. Following the audio presentation, participants completed various behavioral tasks, including comprehension questions, subjective task workload assessments, and reaction time tests which are not discussed in this paper. Finally, participants were given a break before starting the next block. The entire experiment presentation was controlled by custom MATLAB and Psychtoolbox code.

#### Measure of selective attention

Selective attention during listening was measured by the correct identification of color words embedded in the target story, referred to as “hits”. A hit was defined as pressing a designated keyboard button within 2 seconds after the onset of a target word. Any color words in the dual-talker condition that occurred within ±2 seconds of each other in the target and masker were excluded from analyses. Specifically, the relevant measure was the proportion of correct hits out of the total possible, non-overlapping target words (n=25 in the mono-talker condition, n=21 in the dual-talker condition).

### EEG

#### EEG recordings

EEG recordings were obtained in an electrically shielded booth using a BioSemi ActiveTwo system (BioSemi B.V., Netherlands, www.biosemi.com) with a sample rate of 8192 Hz from 64 electrodes following the International 10-20 standard, plus an additional 4 electrode pairs around ears (earlobe, mastoid, posterior to the earlobe and inferior to the mastoids, and superior-anterior to the tragus near the zygomatic arch) for 72 total electrodes. Prior to recording, offsets from all electrodes were measured to be below 20 µV relative to the Common Mode Sense electrode using ActiveView2 software. Data were recorded using Lab Streaming Layer (LSL) software (https://github.com/sccn/labstreaminglayer) in XDF format on a desktop computer running Windows 10. Events were recorded alongside EEG data as a separate trigger channel in the XDF file.

#### EEG Preprocessing

The EEG data were preprocessed in MATLAB 2021b using EEGLAB (Delorme & Makeig, 2004) and custom MATLAB code. Data were first imported from XDF files into EEGLAB using the XDF importer plugin (version 1.19 https://github.com/xdf-modules/xdf-EEGLAB) after which events were extracted from the trigger channel using custom MATLAB code. Given the high number of densely packed event codes (>130,000/subject at an average rate of 1 event every 20ms), events from the 3 different event streams (parallel port and 2 StimTraks) often overlapped in the trigger channel, causing occasional event code errors. We therefore checked all recorded events against expected events and corrected for any missing, extra, or jittered events using custom MATLAB code. We then applied a second set of corrections to adjust for differences in the acoustic signal and the event times. These corrections accounted for the 1.973 ms audio delay due to the HRTF process, a 1 ms audio delay due to the audio travel time from the transducer via the insert earphone tubes, a 1 ms trigger delay in one of the StimTrak channels caused by the TriggerBox hardware, and a 0.541 ms trigger delay to the chirp triggers accounting for a difference between the expected chirp timing from the Cheech process and the chirp onsets.

Data were then pruned to retain only sections of the data from the rest periods and the attention task periods with 10 second buffers on both sides of the selected periods to prevent filtering artifacts. Each remaining section was then DC corrected and filtered using a non-causal 2nd order 0.1 Hz high-pass Butterworth filter using ERPLAB’s filter function to remove slow drifts, and then referenced to electrode Cz. Data were then checked and cleaned for 60 Hz line noise, which was occasionally found in some channels despite the electrically shielded booth. Only channels with detected line noise were cleaned (mean = 4.5 channels ± 7.7 SD, range for a given subject = 0-39 channels) to minimize the effects of the cleaning. This was achieved by first checking for line noise using a frequency tagging approach. Individual channel spectra were calculated across all frequencies. For each frequency, x, the average spectral power of 4 neighboring frequencies except those immediately adjacent (x-2, x-3, x+2, x+3) were subtracted. If the resulting neighbor corrected power at 60 Hz or any of its first 4 harmonics (120, 180, 240, 300 Hz) was greater than 8 standard deviations calculated from the corrected power from all other frequencies, that channel was cleaned of line noise using the Cleanline plugin (https://www.nitrc.org/projects/cleanline). This repeated up to a maximum of 5 times, or until no further line noise was detected.

We used Independent Component Analysis (ICA) to further clean the data of eye blink, eye movement, and heartbeat artifacts following the methodology outlined by Luck (2022). This involved making a copy of the data processed up to this point (ICA copy), down sampling and cleaning the ICA copy, followed by ICA and marking and rejecting artifact components, then transferring the cleaned ICA weight matrix from the ICA copy to the original data. This approach allows for a massive reduction in ICA computation time, while still retaining all the information in the data relevant for decomposing eye and heart artifacts. Specifically, after making a copy for ICA, we down-sampled to 256 Hz to speed ICA processing, and then manually inspected and rejected bad channels (mean=3.67 ± 2.16 SD channels, range=0-9 channels). Following channel rejection, a non-causal 1 to 50 Hz 8th order Butterworth bandpass filter using ERPLAB’s filter function was applied. We then removed sections of data with significant movement and muscle artifacts that can hinder ICA’s ability to isolate eye and heart components, using ERPLAB’s continuous artifact rejection function (mean=126.54 ± 86.85s SD or 3.52% ± 2.36%, range=4.98-298.79s or 0.14-7.96%). The rejection process operated with a 1 s sliding window with a 0.25 s step size and an initial rejection threshold of ±200 µV. In order to retain eye activity for ICA, the rejection threshold was applied against all channels except those mostly likely to contain eye related activations (FP1, FPz, FP2, AF7, AF3, AFz, AF4, AF8). The rejection threshold was inspected and adjusted for each subject to maximize rejecting movement and muscle artifacts while retaining eye blinks (mean=220.83 ±29.18 SD µV, range =200-300 µV). We then used the infomax ICA algorithm after first using Principal Components Analysis (PCA) to reduce the number of channels from an average of 67 to 32 for each subject to further speed ICA. ICA components constituting eye blinks, eye movements, and heart artifacts (when present), were manually selected for rejection (mean=2.79 ± 0.66 SD components, range=2-4 components). The resulting ICA weight matrix was then transferred to the original dataset, after first rejecting the same bad channels in the original dataset as were identified in the ICA dataset.

After the data was cleaned with ICA, we down sampled it to 512 Hz, and applied a non-causal 0.5 to 40 Hz 8th order Butterworth bandpass filter using ERPLAB’s filter function. Data were then re-referenced to the average of the left and right earlobes. In the case that one or both earlobe channels were rejected as bad, the average of the left and right mastoid electrodes was used as reference instead. While re-referencing, activity at channel Cz (the previous reference) was recalculated and placed back into the data. We then used spherical interpolation on any channels rejected.

EEG data were epoched -50 to +500 ms from the first chirp of each word in each condition, with the prestimulus period used as baseline. Each condition had roughly the same number of events (MonoMale = 2417, DualTargetMale = 2392, DualMaskerMale = 2392), except for one subject in which data recording ended early due to computer error (MonoMale = 1249, DualTargetMale = 2000, DualMaskerMale = 2392). Epochs with artifacts were cleaned with a 2-step process: first, a voltage threshold of ±300 µV was applied to remove any epochs with extreme artifacts. Second, any remaining epochs with voltage activity outside of 6 SD for a single channel, or 2 SD across all channels were rejected (MonoMale: mean = 5.01 ±0.83% SD rejected epochs, range = 3.07-6.85%; DualTargetMale: mean = 4.84 ±0.86% SD rejected epochs, range = 3.75-7.02%; DualMaskerMale: mean = 5.12 ±1.1% SD rejected epochs, range = 2.9-8.12%)

#### N1 extraction

N1 component activity was captured as the mean amplitude at electrode Fz between 110 and 160 ms post chirp onset. The electrode site Fz was chosen based on previous research using the Cheech paradigm (Backer et al., 2019) and literature on the scalp regions where auditory evoked responses generally reach peak amplitudes (Kappenman & Luck, 2011). Given the N1 component lacked a clear peak (as is typical during continuous speech presentation) and was usually followed by a sustained negativity (Figures 3, 4), a mean amplitude over the 50 ms interval was used instead of more traditional peak fitting approaches, with the time course chosen based on visual inspection of mono and dual target conditions (Figure 4). Difference scores were calculated by subtracting mean dual target N1 amplitudes from mean dual masker N1 amplitudes.

### Statistical Analysis

All statistical analyses were performed using R version 2023.09.1+494 (Posit Software, PBC) on macOS Sonoma 14.6.1. A paired t-test was performed to evaluate the statistical significance of differences between target and masker N1 amplitudes. A linear multiple regression analysis was conducted to examine how N1 amplitude differences between attended (target) and unattended (masker) speech relates to cognitive and non-audiometric psychoacoustic predictors. The criterion variable was the N1 amplitude difference in the continuous multi-talker spatial attention task. The five predictors included: (1) median reaction time differences between the incongruent and congruent conditions of the Flanker task, (2) median reaction time differences between the incongruent and congruent conditions of the Stroop task, (3) median reaction time differences between TMT-B and TMT-A, (4) letter recall accuracy from 3 and 4 letter trials of the Reading Span test, and (5) average threshold of the Temporal Fine Structure task across 250 and 500 Hz conditions.

The five predictors were entered simultaneously into the multiple regression model to assess their collective effect on the N1 amplitude differences. Prior to the analysis, several diagnostic checks confirmed the suitability of multiple regression. Residual vs. fitted plots and Q-Q plots indicated no significant deviations from normality. Multicollinearity among predictors was assessed using variance inflation factors (VIFs), with values above 5 suggesting problematic multicollinearity (Marcoulides & Raykov, 2019). The highest VIF in our model was 1.43, indicating no substantial effects of multicollinearity. Influential outliers were evaluated using a Cook’s Distance threshold of 0.5. As mentioned previously, one influential outlier was detected and subsequently removed, resulting in a final sample size of N=24. Plots of each predictor against N1 amplitude differences demonstrated no significant deviations from linearity. With a sample size of 24 and five predictors, the subject-per-variable ratio (SPV) was 4.8. A linear regression model requires a minimum SPV of 2 for reliable estimation of regression coefficients, standard errors, and confidence intervals (Austin & Steyerberg, 2015). All variables were z-standardized prior to regression analysis.

Following the multiple regression analyses, dominance analysis was performed to assess the relative importance of predictors in explaining the variance in N1 amplitude differences (Budescu, 1993). This additional analysis was necessary because the regression coefficients can be influenced by even moderate collinearity among predictors, potentially skewing the assessment of each predictor’s contribution (Mizumoto, 2023). Dominance analysis calculates the general dominance weight (GDW) to quantify the variance explained by each predictor. It evaluates the dominance of each predictor by comparing their R² contributions across all possible subset models. Specifically, each predictor is added to all possible models that do not already include it, and the increase in R^2^ is measured. The GDW represents standardized weights of each predictor. Higher GDW values indicate greater explanatory power of a predictor when combined with other predictors.

To validate the predictive accuracy of the regression model, we employed Leave-One-Out Cross Validation (LOOCV). This technique was utilized to ensure the model’s robustness in predicting N1 amplitude differences based on participants’ cognitive and psychophysical performance. In LOOCV, one observation in the dataset (one participant) is used as a validation set, while the remaining datasets (N-1) are used as training data to predict the value for the omitted observation. This iterative process is repeated N times, once for each observation. For each iteration, the model’s prediction error is calculated, and these errors are aggregated to compute the root-mean-squared error (RMSE), which provides a measure of the average magnitude of prediction errors. This method provides a nearly unbiased estimate of the model’s predictive performance, particularly beneficial for smaller datasets.

A separate multiple regression analysis was conducted using the same five predictors to examine their ability to predict the proportion of color word hits in the continuous multi-talker spatial attention task. The predictors were entered simultaneously into the regression model to assess their combined impact on speech perception performance in noise. Prior to the analysis, the appropriateness of using multiple regression was verified through several diagnostic checks. Although the residuals vs. fitted plots and Q-Q plots indicated some deviations from normality, no transformation of the response variable significantly improved the distribution of the residuals or altered the results. Since the same predictors were used, the multicollinearity checks from the first model remained unchanged. The influential outlier observed in the first model was also removed in this analysis (N=24). Plots of each predictor against N1 amplitude differences showed no significant deviations from linearity. All variables were z-standardized prior to regression analysis.

## RESULTS

### Cognitive and Psychoacoustic Predictors

Figure 2 presents box plots illustrating the distribution of scores across the five cognitive and psychoacoustic tasks administered to participants, highlighting the variability in performance. While some plots reveal visually apparent outliers, these data points did not influence the predictive performance of the models and were therefore retained in the analysis.

**Figure 2:**
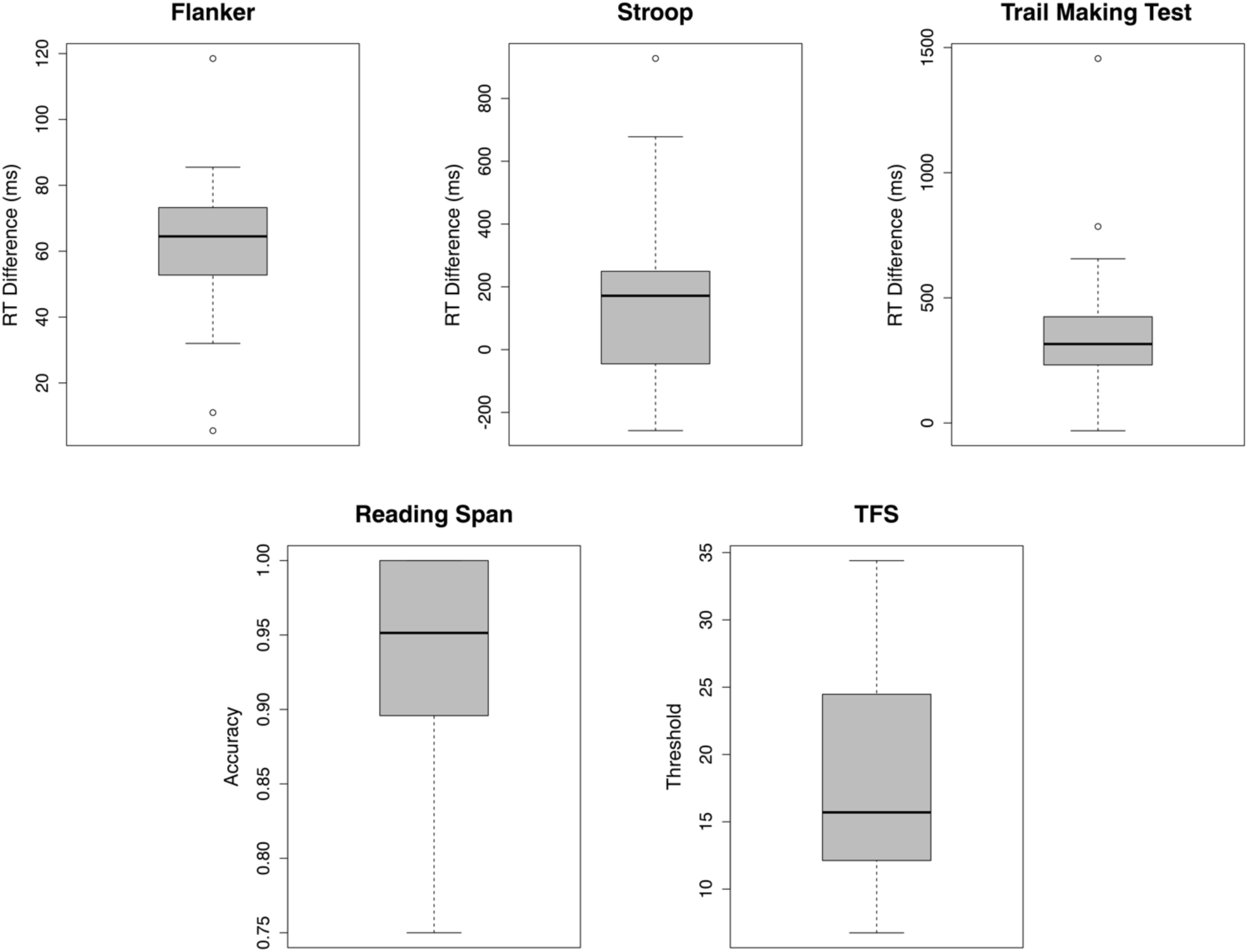
Box plots illustrating the distribution of performance scores across five cognitive and psychoacoustic predictors (N=24). Larger RT difference scores (Flanker, Stroop, and Trail Making) indicate greater response (cognitive) interference in each domain (i.e., slower RTs for the incongruent/alternating trials than the congruent/sequential trials).

### N1 Across Conditions

The grand average waveforms in Figure 3A reveal a significant difference in N1 amplitudes between the target and masker story conditions. The N1 amplitude was significantly larger for the target story condition (M = -0.142 µV, SD = 0.350 µV) compared to the masker story condition (M = 0.113 µV, SD = 0.198 µV). A paired t-test confirmed this difference as statistically significant (t(23) = -3.664, p = 0.001). Figure 3B displays the difference wave, created by subtracting the ERP waveform of the masker from that of the target. Negative N1 amplitude differences indicate a stronger N1 response to target speech compared to masker speech, while positive differences suggest a weaker N1 response to target speech. The average N1 amplitude difference between the target and masker speech was -0.254 ± 0.340 µV.

**Figure 3:**
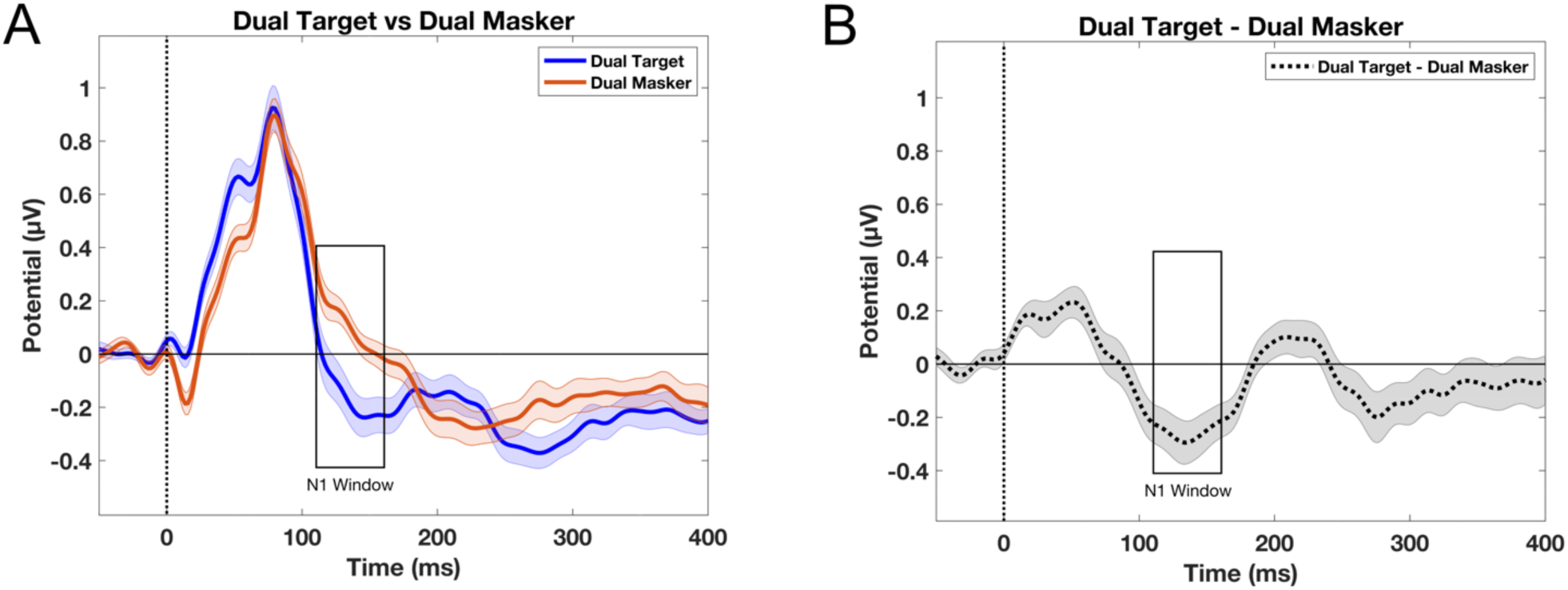
Attentional modulation of speech-evoked neural activity in dual-talker conditions at electrode Fz. Error bars indicate standard error. A) Grand average ERP waveforms of the target and masker speech. B) Difference waveforms between target and masker stories.

After analyzing the grand average waveforms for the target and masker story conditions, we compared them to a baseline mono condition, in which there was only one speaker for participants to attend to. This comparison was conducted to determine whether selective attention in a dual-talker scenario primarily boosts the neural response to the target speech, suppresses the response to competing speech, or involves both processes. The N1 amplitude did not differ significantly between the (dual) target and mono story conditions, as indicated by a Bonferroni-corrected paired t-test (t(23) = -0.790, p = 0.438; see Figures 4A and 4B). However, the N1 amplitude for the masker story condition was significantly smaller than that of the mono story condition (t(23) = -2.798, p = 0.010), as shown in Figures 4C and 4D.

**Figure 4:**
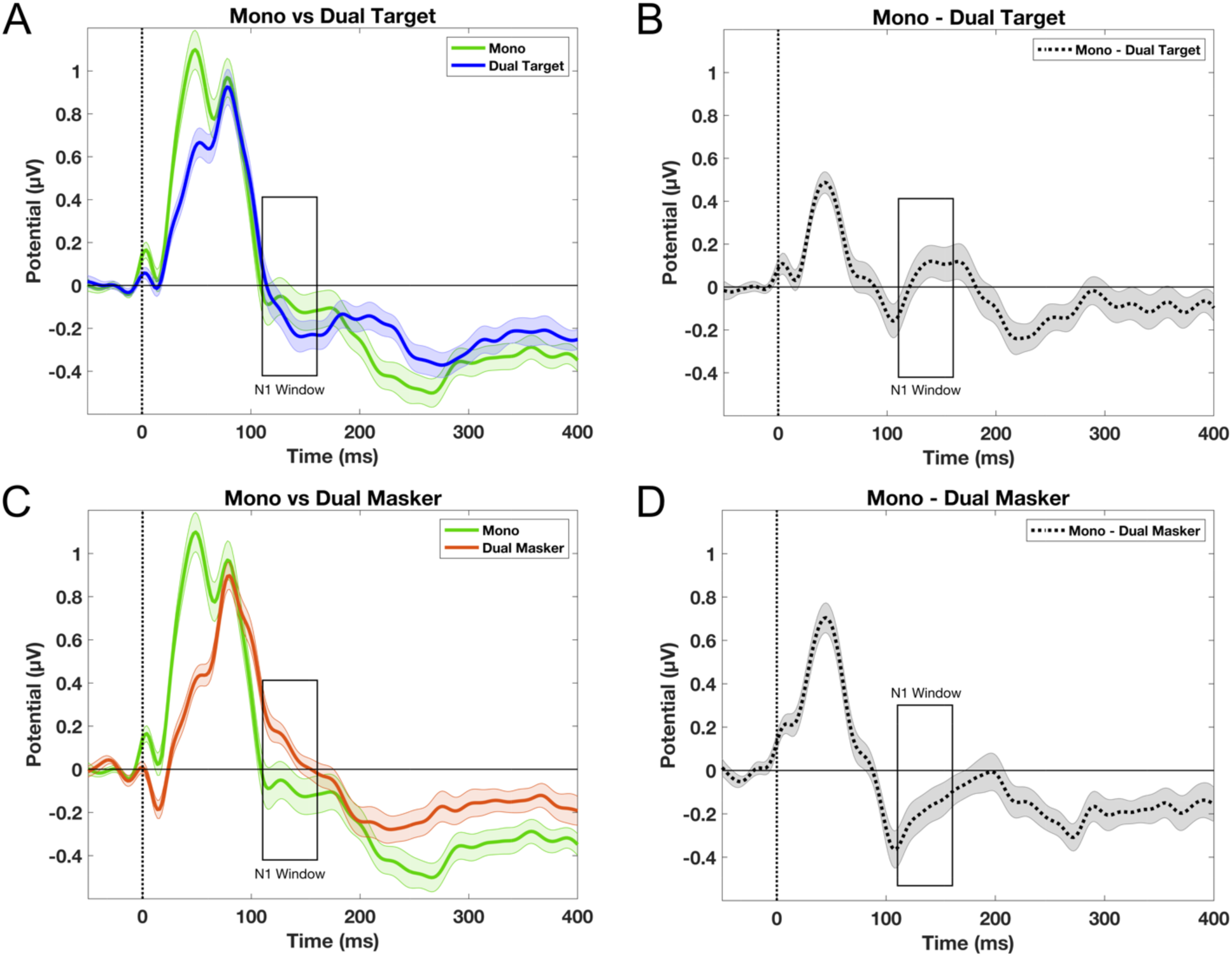
Attentional modulation of speech-evoked neural activity comparing mono and dual-talker conditions at electrode Fz. Error bars indicate standard error. A) Grand average ERP waveforms comparing the dual-talker target condition to the mono condition. B) Difference waveforms (mono - target), where positive values indicate larger N1 amplitudes in the dual-talker target condition than in the mono condition. C) Grand average ERP waveforms comparing the dual-talker masker condition to the mono condition. D) Difference waveforms (mono - masker speech), where negative values indicate smaller N1 amplitudes in the dual-talker masker condition compared to the mono

### Predicting N1 Amplitude Differences

We used multiple regression analysis to assess whether performance on cognitive and psychoacoustic tasks could predict N1 amplitude differences between target and masker narratives (i.e., dual-talker condition only). As shown in Table 1, the multiple regression model demonstrated a strong fit.

**Table 1:** Results of the multiple regression analysis. Criterion variable: N1 amplitude difference between target and masker stories in the continuous multi-talker spatial attention task. Predictors: performance on (1) Flanker, (2) Stroop, (3) Trail Making Test, (4) TFS, and (5) Reading Span. All variables were z standardized. GDW (General Dominance Weight) indicates the relative importance of each predictor in the model. Overall adjusted R^2^ = 0.467, p = 0.005.

| | $\beta$ | SE | t | p | GDW |
| --- | --- | --- | --- | --- | --- |
| Intercept | 0.000 | 0.149 | 0.000 | 1.000 |  |
| Flanker | 0.047 | 0.168 | 0.277 | 0.784 | 0.009 |
| Stroop | 0.598 | 0.178 | 3.370 | 0.003 | 0.284 |
| Trail Making Test | -0.942 | 0.182 | -0.518 | 0.611 | 0.044 |
| Reading Span | 0.649 | 0.167 | 3.894 | 0.001 | 0.477 |
| TFS | -0.359 | 0.182 | -1.971 | 0.064 | 0.185 |

The N1 amplitude difference between target and masker narratives was significantly positively associated with Reading Span Accuracy on three and four-letter trials. This indicates that better working memory performance was associated with reduced differentiation in auditory cortical responses (i.e., closer to 0 or more positive N1 amplitude difference between target and masker). In contrast, the N1 amplitude difference demonstrated the opposite relationship with Stroop Task reaction time differences between incongruent and congruent trials. Specifically, stronger behavioral inhibition abilities, in the form of reduced response interference (i.e., *smaller* Stroop RT difference values), were associated with more robust attentional effects in the auditory cortex (i.e., larger target-compared to masker-evoked N1 responses).

TFS threshold and reaction time differences between incongruent and congruent trials on the Flanker and Trail Making Test were not statistically significant predictors. However, there was a trend suggesting that better TFS ability (indicated by lower thresholds) was associated with smaller N1 amplitude differences. Figure 5 illustrates the relationships between each individual predictor and N1 difference amplitudes. Additionally, Figure 6 provides further insights by depicting the confidence intervals for each predictor’s coefficients.

**Figure 5:**
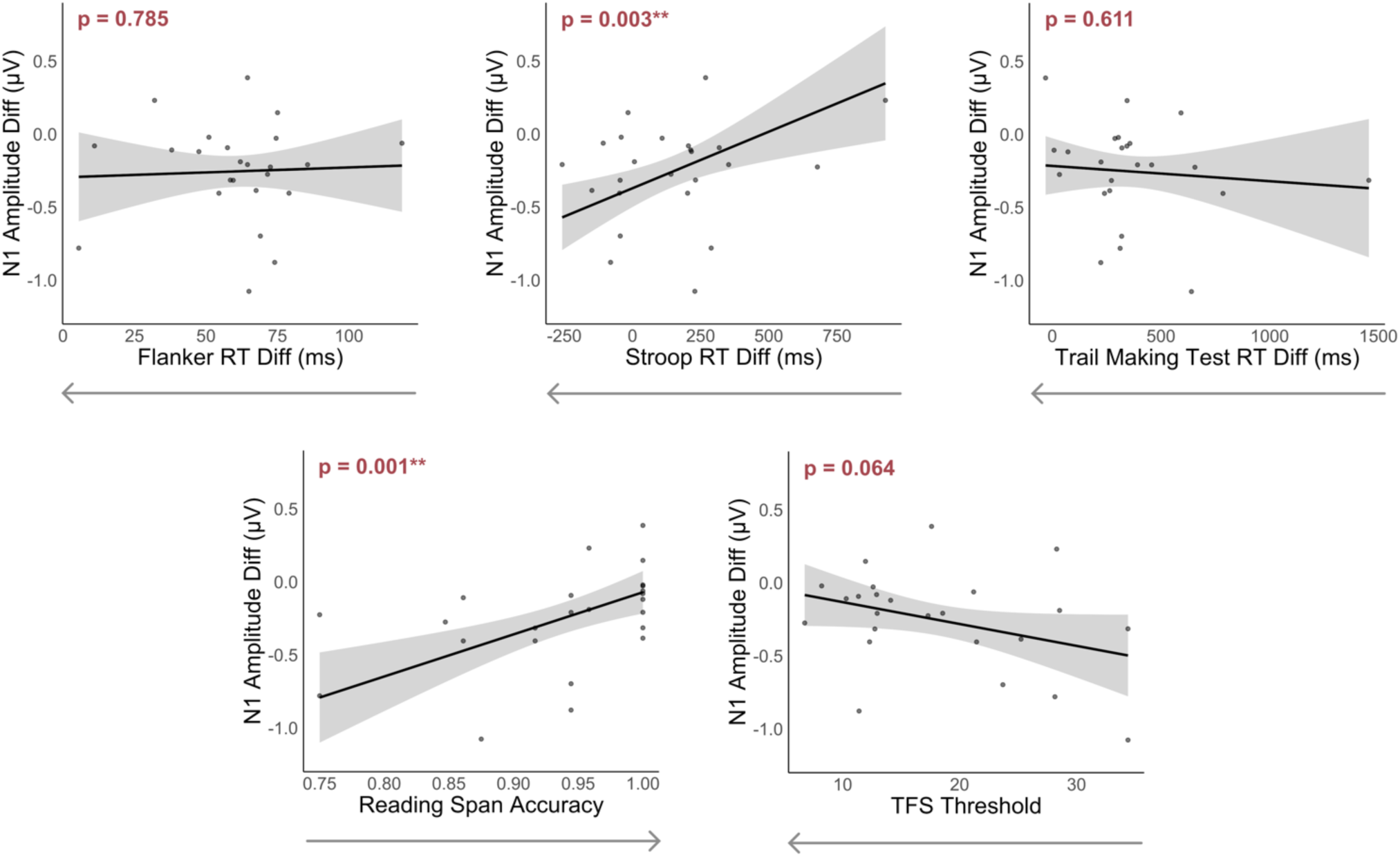
Relationship between each predictor and the N1 amplitude difference. All other predictors are held constant. Shaded regions represent 95% confidence intervals. N1 amplitude differences are calculated as the N1 amplitude for the target minus that of the masker; lower values on the y-axis correspond to larger differences in N1 amplitude between the target and masker streams. Arrows on the x-axis denote better performance on the cognitive and psychoacoustic tests.

**Figure 6:**
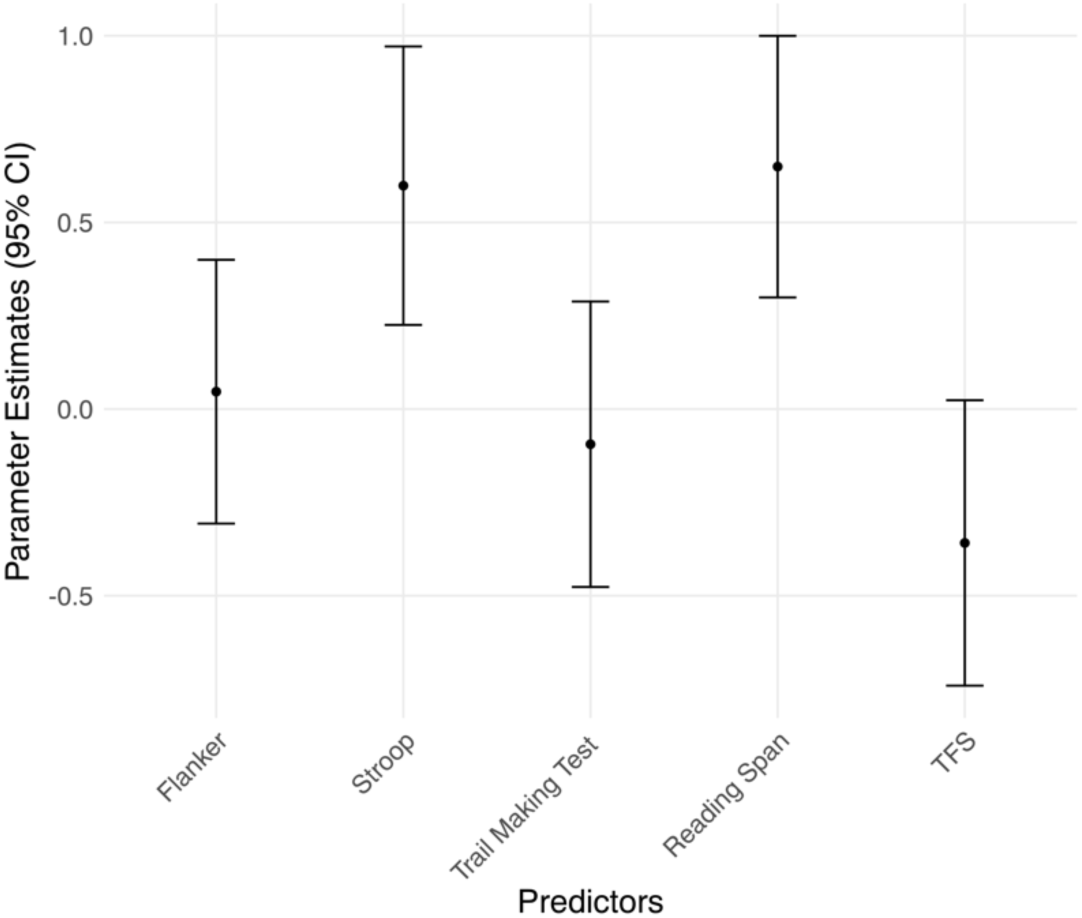
95% confidence intervals for the regression coefficients of each predictor.

Leave-One-Out Cross Validation (LOOCV) was used to validate the model fit and assess its predictive performance. The scatter plot in Figure 7 illustrates the relationship between the actual N1 amplitude differences (x-axis) and the predicted N1 amplitude differences (y-axis) generated from the LOOCV procedure. Each point on the plot represents an individual observation used as the validation set in a single iteration of LOOCV. The diagonal line indicates perfect prediction where predicted values match the actual values. Points closely aligned with this line indicate accurate predictions, while points further away indicate prediction errors. The LOOCV analysis resulted in a root-mean-square error (RMSE) of 0.215 microvolts, reflecting the average magnitude of prediction errors for the N1 amplitude differences.

**Figure 7:**
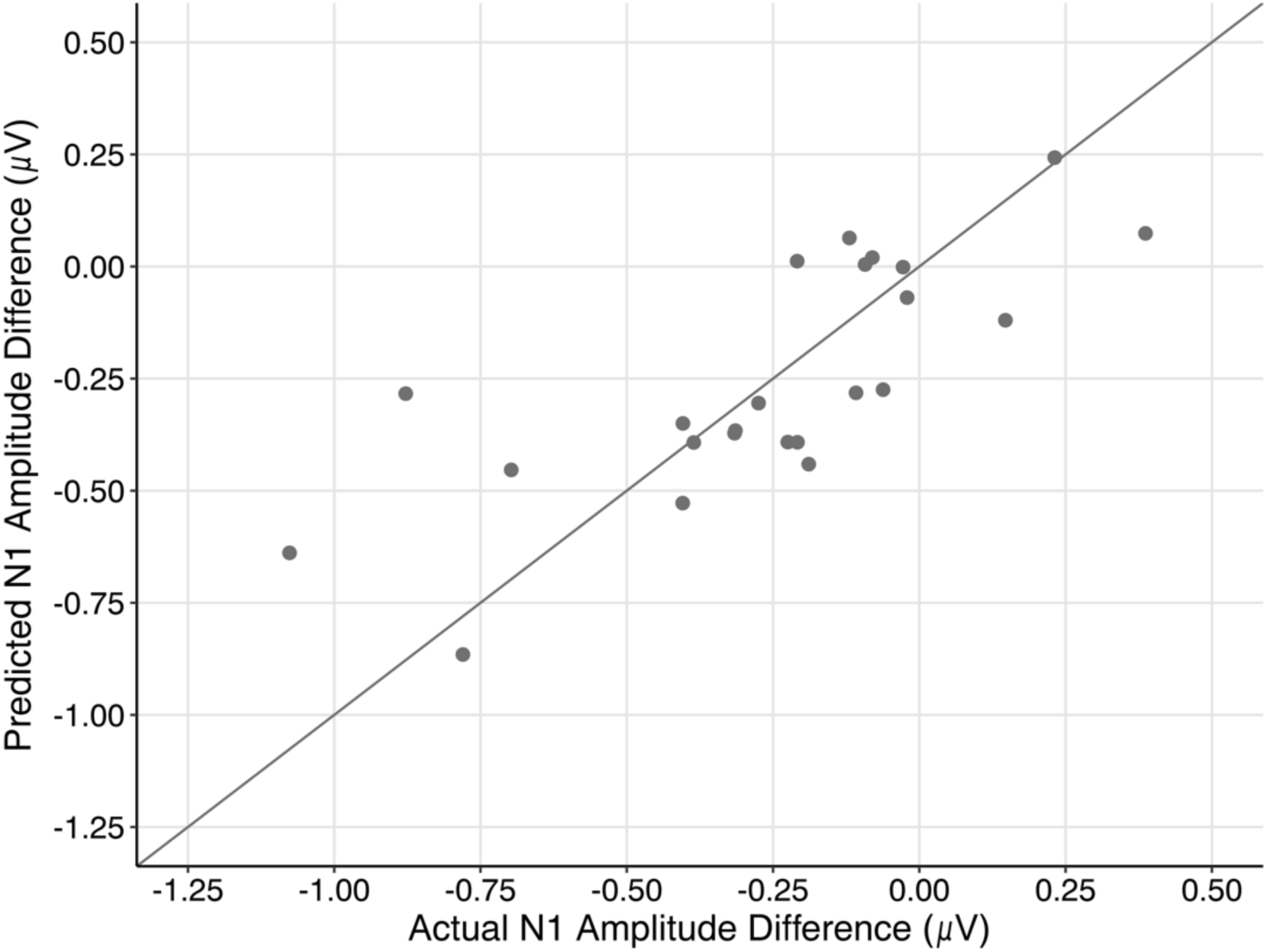
Cognitive and psychoacoustic abilities predict attentional modulation of speech representations. Results of the LOOCV. The graph shows the predicted and actual N1 amplitude differences for each participant.

As previously mentioned, none of the six predictors exhibited strong collinearity, as indicated by low Variance Inflation Factors (VIFs), suggesting that the model and its parameter variances should be stable. Nonetheless, the predictors were mildly correlated (see Table 2). In such situations, relying solely on regression coefficients to assess predictor importance can be misleading. To address this, we employed the "dominance analysis" method (Budescu, 1993) to evaluate the relative importance of each predictor in the regression model. The results revealed that Reading Span had the highest general dominance weight (GDW) across both models, followed by Stroop, indicating that working memory and response inhibition play significant roles in driving neural encoding differences between target and masker speech (see Table 1).

**Table 2:** Pairwise Pearson correlation coefficients of the performance on cognitive tasks.

|  | Flanker | Stroop | Trail Making | Reading Span | TFS |
| --- | --- | --- | --- | --- | --- |
| Flanker | $r = 1.000$ | $r = -0.369$<br>$p = 0.076$ | $r = 0.065$<br>$p = 0.763$ | $r = 0.245$<br>$p = 0.248$ | $r = -0.127$<br>$p = 0.555$ |
| Stroop | | $r = 1.000$ | $r = 0.156$<br>$p = 0.467$ | $r = -0.393$<br>$p = 0.057$ | $r = 0.196$<br>$p = 0.358$ |
| Trail Making | | | $r = 1.000$ | $r = -0.023$<br>$p = 0.917$ | $r = 0.520$<br>$p = 0.009$ |
| Reading Span | | | | $r = 1.000$ | $r = -0.075$<br>$p = 0.727$ |
| TFS | | | | | $r = 1.000$ |

Interestingly, the strongest correlation observed among the predictors was between TFS thresholds and performance on the Trail Making Test. A positive correlation was found, suggesting that individuals with lower (better) TFS thresholds tended to exhibit smaller reaction time differences between the A and B conditions on the Trail Making Test, indicating better executive function (i.e., less cognitive susceptibility to task switching delays in the TMT-B).

### Predicting Behavioral Performance of Speech Perception in Noise

The earlier models explored the relationships between neural responses and performance metrics obtained separately from the EEG session. However, during the EEG session, participants also engaged in behavioral tasks that approximate real-world listening abilities. To further explore these relationships, we conducted an additional analysis using the same predictors to predict the proportion of color word hits, defined as the proportion of correctly identified color words in the target story within two seconds of their onset (i.e., excluding overlapping color words across the story pairs; Figure 8). This measure was selected because it directly reflects the participants’ ability to focus on the target speech and respond promptly, making it a relevant and sensitive measure of real-world listening performance. We aimed to assess how well the observed relationships of the predictors with neural activity translate to behavioral outcomes. Based on this multiple regression model, none of the predictors were correlated with the proportion of color word hits. This finding suggests that while cognitive and psychoacoustic abilities can accurately predict neural activity during SiN perception, they do not appear to predict SiN behavioral performance in this cohort. The relationship between each predictor and the proportion of color word hits is illustrated in Figure 9.

**Figure 8:**
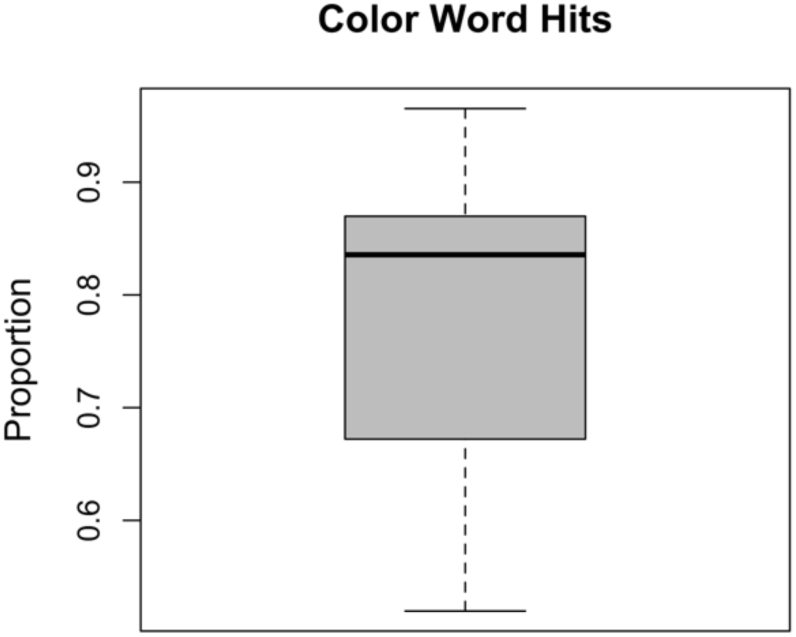
Box plot illustrating the distribution of proportion of color word hits in the continuous multi-talker spatial attention task.

**Table 3:** Results of the second multiple regression analysis. This model is using the same five cognitive and psychoacoustic predictors to predict the proportion of color word hits in the continuous multi-talker spatial attention task. All variables were z standardized. Overall R^2^ = -0.144, p = 0.828.

| | $\beta$ | SE | t | p | GDW |
| --- | --- | --- | --- | --- | --- |
| Intercept | 0.000 | 0.218 | 0.000 | 1.000 | 0.000 |
| Flanker | -0.020 | 0.246 | -0.082 | 0.935 | -0.020 |
| Stroop | 0.070 | 0.260 | 0.269 | 0.791 | 0.070 |
| Trail Making Test | -0.348 | 0.267 | -1.305 | 0.208 | -0.348 |
| Reading Span | 0.135 | 0.244 | 0.551 | 0.589 | 0.135 |
| TFS | 0.230 | 0.267 | 0.861 | 0.400 | 0.230 |

**Figure 9:**
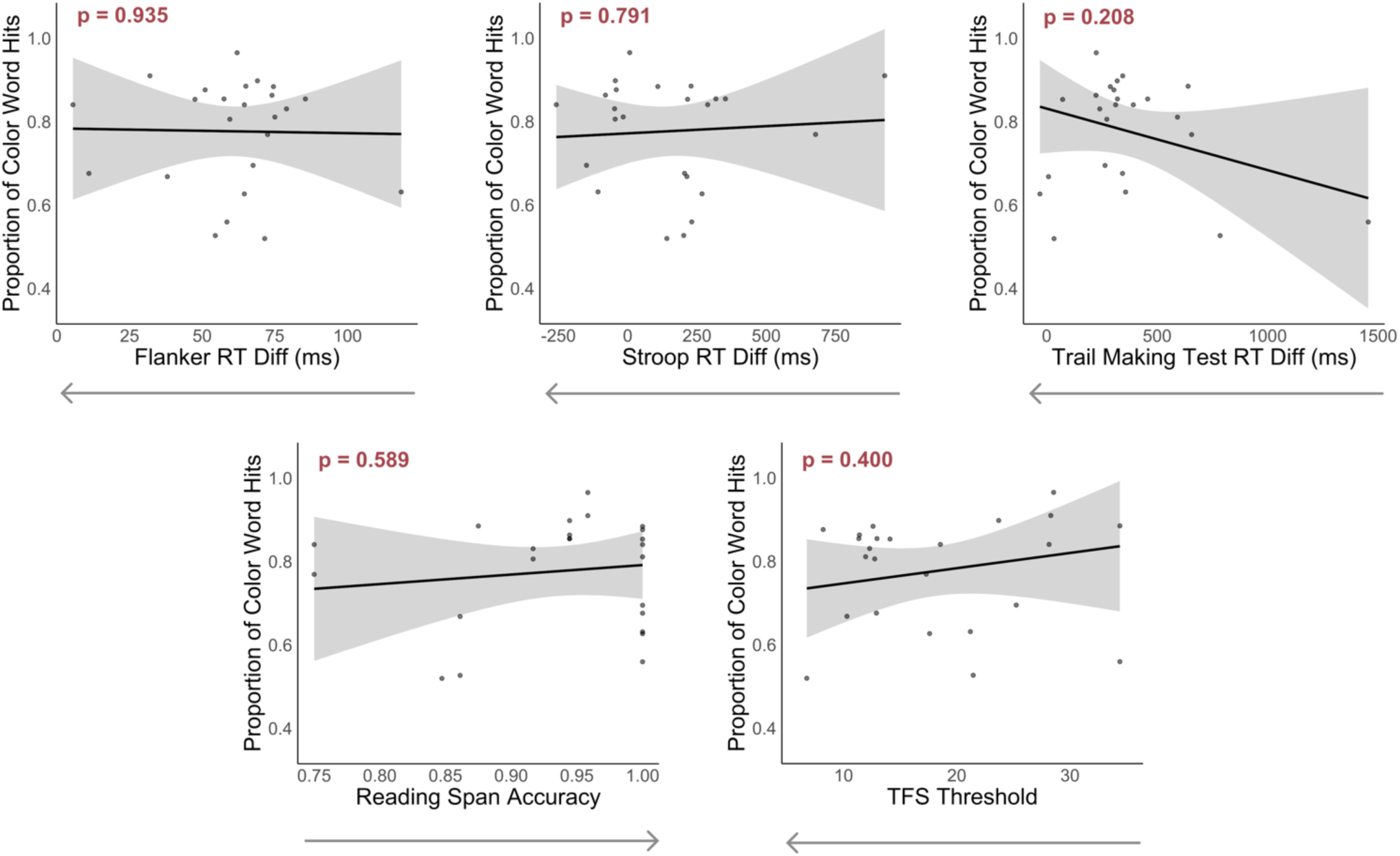
Scatterplots showing the relationship between each predictor and the proportion of color word hits. All other predictors held constant. The shaded regions represent 95% confidence intervals of the proportion of color word hits.

### Relationship Between N1 Amplitude Differences and Proportion of Color Word Hits

To gain further insight into the relationship between neural processing and speech perception, we performed a simple linear regression between the N1 amplitude differences and the proportion of color words that participants correctly identified. N1 amplitude differences between target and masker speech were not significantly associated with this ecologically valid SiN performance metric (Figure 10).

**Figure 10:**
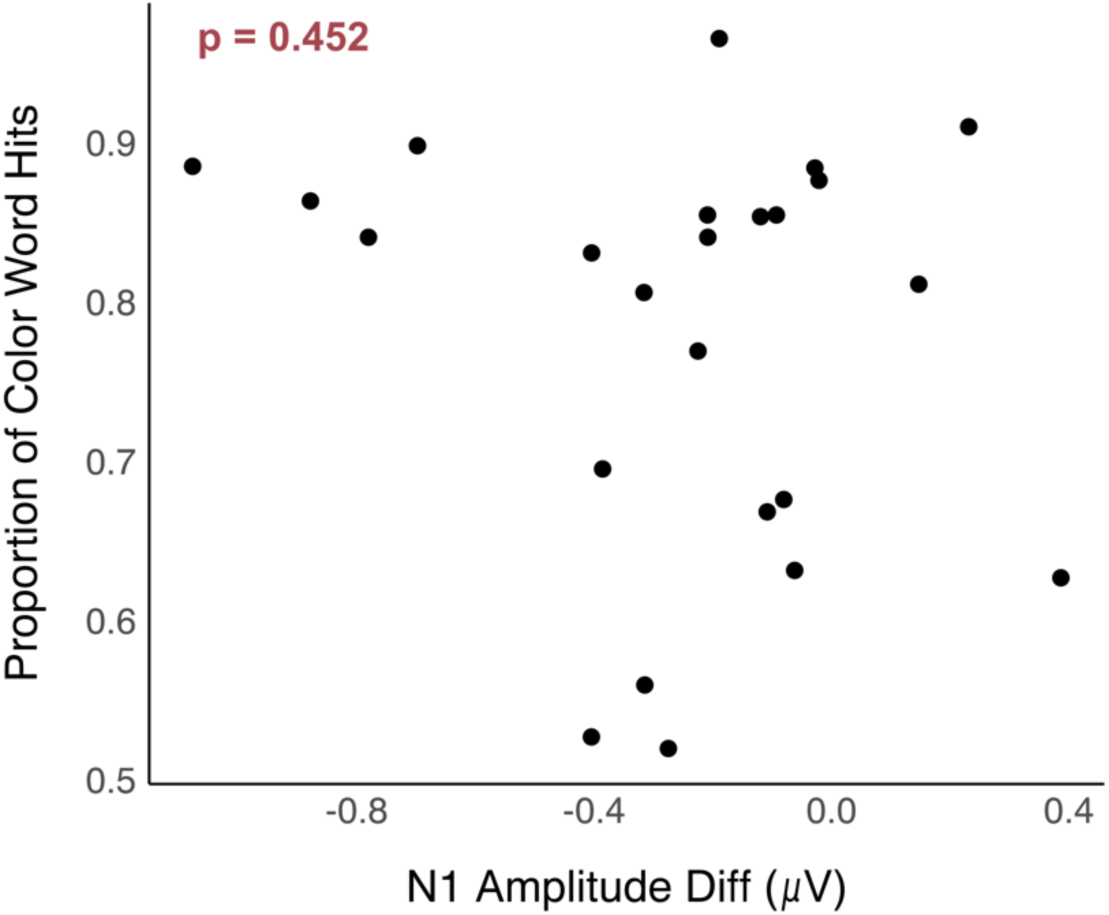
Scatter plot showing the relationship between the proportion of color word hits and N1 amplitude differences.

## DISCUSSION

In this study, we aimed to identify the key predictors of neural processing within the auditory cortex (specifically N1 amplitudes) associated with differentiating target from masker speech in a multi-talker spatial attention task among normal hearing listeners. For clarity, we adhered to well-established interpretations of each cognitive predictor, emphasizing the constructs most relevant to our dynamic speech-in-noise (SiN) EEG task. Specifically, Reading Span gauged working memory abilities, the Stroop task measured linguistic interference and inhibitory control, the Flanker task assessed selective attention and stimulus inhibition, and the Trail Making Test evaluated broader executive functions, including cognitive-attentional switching and processing speed. Additionally, Temporal Fine Structure (TFS), a psychoacoustic measure, was included as it reflects sound localization abilities.

### Predicting N1 Amplitudes

Our multiple regression model, which included five cognitive and non-audiometric psychophysical predictors, demonstrated that working memory performance and inhibitory control significantly contributed to the variance in N1 amplitude differences between target and masker talkers. TFS also showed a trend toward further explaining additional variance. These findings demonstrate that, even among audiometrically normal listeners, SiN processing varies based on domain-general abilities. This highlights the complexity of neural mechanisms involved in comprehending speech in noisy environments, which span from basic auditory processing to higher-level cognitive functions.

Our multivariate approach further examines the relationship between each predictor and N1 amplitude differences between target and masker stories. Notably, accuracy on the Reading Span task for three- and four-letter trials was a significant predictor, emphasizing the role of working memory in neural encoding processes during challenging listening scenarios. Interestingly, higher accuracy on the Reading Span task was associated with *smaller* N1 amplitude differences, indicating that individuals with better working memory exhibit more efficient neural processing (i.e., smaller, or more positive N1 response differences; closer similarity between target and masker responses). Previous research has shown that individuals with higher cognitive performance use fewer neural resources during demanding tasks, a concept known as neural efficiency (Haier et al., 1988). As individuals listen to spoken language, they apply their knowledge of syntax, semantics, real-world context, and conversational rules to predict upcoming content (Akeroyd, 2008). This predictive processing helps in segmenting the continuous acoustic stream into words and making inferences more efficiently. Therefore, individuals with greater working memory capacity may be more efficient at these predictive processes, reducing the need for pronounced neural differentiation between target and masker speech, as reflected in smaller N1 amplitude differences.

Interestingly, in our pilot testing, the total Reading Span accuracy of all trials (3 to 7 sentence-letter pairs) was not a strong predictor of N1 amplitude differences between target and masker speech. This finding suggests that accuracy on three- and four-letter trials may be a more informative measure of working memory capacity related to neural processes in SiN perception. One possible explanation is that tasks involving fewer items (three to four letters) fall within the working memory capacity limit (Cowan, 2001, 2015). This smaller capacity measure might be more sensitive to individual differences in neural efficiency and attentional control during auditory attention tasks. When the Reading Span exceeds this capacity (five to seven letters), performance may be influenced by additional factors such as strategies, chunking, or long-term memory processes, which may not directly reflect the working memory capacity relevant to real-time speech processing in noise. Consequently, focusing on moderate working memory capacities may provide a more accurate representation of the cognitive resources used during complex listening tasks, leading to better predictions of neural activity differences in multi-talker auditory environments.

In our study, better performance on the Stroop task—indicated by smaller reaction time differences between incongruent and congruent conditions— was associated with larger N1 amplitude differences between target and masker stories. The Stroop task is designed to assess linguistic interference and inhibitory control (Stroop, 1935; Troyer et al., 2006). This task requires participants to inhibit the automatic tendency to read the word and instead name the ink color, demonstrating their ability to suppress irrelevant information. This skill is likely transferable to complex listening environments, where participants must focus on target speech while inhibiting masker speech. Individuals with stronger inhibitory control may be better at suppressing neural representations of distracting information, allowing for enhanced focus on the target speech.

To further this point, we compared the average N1 amplitudes of the dual-talker conditions to the single-talker (mono) condition within the continuous multi-talker spatial attention task. The average N1 amplitude for the masker stream was smaller than the single-talker condition. In the single-talker condition, there is no need for inhibitory control as there are no competing stimuli. However, in a dual-talker scenario, effective inhibition of irrelevant speech is crucial. Thus, these results indicate individuals with better inhibitory control have significantly inhibited N1 responses to masker streams. This supports previous findings showing that the N1 is sensitive to attentional processes (Luck, 1995), with larger N1 amplitude differences reflecting greater attentional resource allocation for the target speech (Ding & Simon, 2012; Hillyard et al., 1973; Kerlin et al., 2010).

Temporal Fine Structure (TFS) thresholds did not significantly predict N1 amplitude differences. Nevertheless, as shown in Figure 5, there is a notable trend suggesting that better TFS thresholds are associated with smaller N1 amplitude differences between target and masker speech. TFS processing starts in the peripheral auditory system, beginning with basilar membrane filters in the cochlea and extending through the brainstem (Borjigin, 2023), and is clearly important for auditory object formation and individuation in cluttered scenes (Moore, 2008). During active listening, attentional mechanisms often operate at the level of perceptual objects, including at the level of the N1 (Hillyard et al., 1973). Thus, better TFS may support more precise, distinct speech representations (target vs masker) at the cortical level, which then would require less attentional gain modulation to distinguish them for further linguistic and other processing. This highlights the need for further research to better understand the relationship between peripheral auditory processing and cortical neural responses, particularly in challenging listening scenarios.

The Trail Making Test (TMT) reaction times were not significant predictors of N1 amplitude differences in our study. Although cognitive flexibility may be important for switching between tasks and managing multiple streams of information, it may not directly impact the neural processes involved in sustaining attention to a target speaker amidst background noise, even with dynamic attentional switching as in our continuous multi-talker spatial attention task. Additionally, the TMT involves significant visual-motor coordination and task-switching, engaging different neural pathways and cognitive resources than those required for auditory attention and speech processing. As a result, the TMT’s emphasis on visual search and motor responses may not effectively translate to tasks involving auditory processing.

Flanker test reaction times were not significant predictors of N1 amplitude differences in our study. Although the Flanker test is a well-established measure of selective attention in (visual) space, it may not fully capture the neural processes required to differentiate between target and masker speech in a noisy environment. The Flanker, like the Stroop, also assesses inhibitory control, but does so in a different context. The Stroop test requires participants to inhibit a more automatic linguistic response (reading the word) in favor of a less automatic one (naming the ink color), directly engaging cognitive mechanisms related to speech and language processing. This linguistic interference is evidently relevant to the challenges of SiN perception, especially when maskers are meaningful speech rather than non-linguistic sounds like steady-state noise, which does not trigger lexical activity (Lu et al., 2016; Schneider et al., 2022). In contrast, the Flanker test primarily involves visuo-spatial interference, where participants must inhibit responses to maskers visually adjacent to the target stimulus. Although our continuous multi-talker spatial attention task relies on inhibiting a speech stream that is spatially adjacent to the target, it is interesting to observe a stronger relationship with Stroop performance than with Flanker performance. This suggests that in individuals with normal hearing performing a realistic SiN task, linguistic interference is more impactful than (visuo-)spatial interference. Future work might determine whether this asymmetry also holds when listeners have hearing loss and, consequently, greater challenges with spatial masking.

### Implications for the N1

Our findings suggest a complex interaction between cognitive abilities and auditory processing, updating our understanding of the N1 component. Traditionally considered a marker of basic auditory processing and attention, the N1 also appears to be influenced by higher-order cognitive functions such as inhibition and working memory, particularly in challenging listening scenarios. Extensive research suggests that when individuals attend to specific auditory stimuli, the N1 amplitude is typically enhanced, indicating a greater neural response to the attended sounds (Hillyard et al., 1973; Kerlin et al., 2010; Picton & Hillyard, 1974; Stapells, 2009). Consistent with this, better response inhibition, as measured by the Stroop task, is associated with larger N1 amplitude differences. However, our findings from the Reading Span test indicate that this relationship is more complicated; higher working memory capacity is associated with *smaller* N1 amplitude differences between attended and unattended streams. This suggests that neural processing during speech perception is determined by compensations and compromises between top-down cognitive influences.

### Predicting Behavioral Performance of Speech Perception in Noise

In addition to analyzing the direct relationship between cognitive performance and neural processing (N1 amplitude differences), we also assessed how each cognitive predictor relates to ecologically relevant listening behaviors during the continuous multi-talker spatial attention task, conducted simultaneously while EEG was recorded. Specifically, we tested the relationship of each cognitive predictor and the proportion of correctly identified color words in the target story. Interestingly, none of the cognitive and psychoacoustic predictors showed a significant correlation with this behavioral measure.

Our cohort consisted of young, highly educated individuals, with all participants having completed at least 13 years of education. The well-educated status of these participants likely equips them with advanced compensatory mechanisms that enable strong performance in speech perception tasks, despite differences in cognitive abilities. As shown in Figure 8, participants achieved a high proportion of color word hits, indicating strong overall performance. For those with lower cognitive task performance, their educational background and overall health may have provided them with effective strategies for maintaining high performance in challenging listening tasks. These strategies might include better use of contextual cues, more effective allocation of attentional resources, and superior problem-solving skills, allowing them to compensate for cognitive limitations. Consequently, the homogeneity in educational background might mask differences in cognitive performance, resulting in consistently strong performance in the continuous multi-talker spatial attention task, regardless of cognitive predictor scores.

Although the relationships between cognitive tests and listening performance *were not* statistically significant, it is noteworthy that some correlations with N1 amplitudes *were* significant. This indicates that neural measures, such as N1 amplitude differences, may serve as more sensitive or direct indicators of auditory processing efficiency and cognitive resource allocation during SiN perception than behavioral measures alone. That is, neural measures can capture subtle aspects of cognitive and auditory processing that behavioral measures, like color word detection, might fail to detect. No single behavioral speech test, such as color word detection, can readily parse the many underlying mechanisms that might contribute to it, particularly, as noted above, when “less is more” (efficiency) or when compensatory mechanisms restore behavioral performance in the presence of an underlying deficit. In these cases, neural metrics can add substantially to our understanding which might not be fully captured by behavioral data alone. Finally, N1 amplitude differences were also not associated with the proportion of color word hits in the target story. This suggests that the behavioral measure likely reflects the influence of multiple underlying neural mechanisms, not exclusively N1 amplitude attentional modulations.

### Relationship between TFS and Trail Making Test

As illustrated in Table 2, there is a notable correlation between TFS thresholds and reaction time differences on the Trail Making Test, indicating that individuals with better TFS processing abilities may also demonstrate greater cognitive flexibility and executive functioning. This link aligns with previous findings by Füllgrabe et al. (2015), who reported similar associations. The underlying mechanisms associating TFS processing and executive function are not well understood. Further research is needed to understand the neural processes that may link TFS and executive function.

## CONCLUSION

Overall, our findings highlight the complexity of the relationships between auditory processing, cognitive abilities, and attentional gains in speech perception under noisy conditions. We demonstrate that neural measures reflect impactful inter-individual differences that are not fully captured by solely traditional assessments of audiometric hearing and behavioral performance. These findings have significant implications for improving hearing loss diagnostics, which currently rely primarily on ear health assessments. By incorporating cognitive abilities and neural activity measures, we can obtain a more comprehensive understanding of speech perception, even among audiometrically normal listeners. Having established the predictive value of these tasks for neural processing of SiN, future work might further clarify *how* each of these reflects differences in underlying cognitive mechanisms across individuals. Understanding how the brain processes auditory information in noisy environments—and how differences in cognition influence this process—could lead to earlier detection of hearing loss, improved strategies for enhancing speech perception in daily life, and ultimately, a significant improvement in the quality of life for individuals facing hearing challenges.

## Acknowledgements

This research was supported by the Department of Defense (DoD) and the Child Family Fund for the Center for Mind & Brain. The authors would like to thank members of the University of California, Davis Health audiology and clinical research team for performing initial hearing evaluations of our participants: Dr. Robert Ivory, Au.D., Dr. Mackenzie Quinn, Au.D., Dr. Rachel Krager, Au.D., Dr. Steven Zurawski, Au.D., Dr. Austin Childers, Au.D., Dr. Kimberly Smith, Au.D., Randev Sandhu, and Angela Beliveau. We extend our thanks to Cathleen Chan and Jillian McKie for their assistance in data collection. We are also grateful to Elyse Ehlert and Tiana Smith for their efforts in participant recruitment. We would like to acknowledge Sophie Burstein, Alicia Dye, Reina Itakura, Zachary McNaughton, Ferdous Rahimi, Tyler Statema, Audrey Vargas, and Nina Wade for their insightful discussions and constructive feedback. Special thanks to Dr. Chhayakant Patro for providing us with the computerized version of the Trail Making Test. Finally, we sincerely thank the participants of this research study without whom none of this work would be possible. This research is published in remembrance of our team member, lab mate, and friend, Karim Abou Najm.

## Conflicts of interest

Dr. Lee M. Miller is an inventor on intellectual property related to chirped-speech (Cheech) owned by the Regents of the University of California and not presently licensed.

## Author contributions

D.C.C, K.M, H.B, D.S, and L.M.M designed research; S.S, B.B, and S.D performed research; D.C.C, K.M, B.B, and L.M.M contributed analytic tools; S.S and D.C.C analyzed data; S.S, D.C.C, K.M, B.B, S.D, D.S, and L.M.M wrote and edited the paper.

## References

1. Akeroyd, M. A. (2008). Are individual differences in speech reception related to individual differences in cognitive ability? A survey of twenty experimental studies with normal and hearing-impaired adults. International Journal of Audiology, 47(sup2), S53–S71. 10.1080/14992020802301142

2. Alain, C., & Arnott, S. R. (2000). Selectively attending to auditory objects. Frontiers in Bioscience: A Journal and Virtual Library, 5, D202–212. 10.2741/alain

3. Armstrong, C., Thresh, L., Murphy, D., & Kearney, G. (2018). A Perceptual Evaluation of Individual and Non-Individual HRTFs: A Case Study of the SADIE II Database. Applied Sciences, 8(11), Article 11. 10.3390/app8112029

4. Austin, P. C., & Steyerberg, E. W. (2015). The number of subjects per variable required in linear regression analyses. Journal of Clinical Epidemiology, 68(6), 627–636. 10.1016/j.jclinepi.2014.12.014

5. Backer, K. C., Kessler, A. S., Lawyer, L. A., Corina, D. P., & Miller, L. M. (2019). A novel EEG paradigm to simultaneously and rapidly assess the functioning of auditory and visual pathways. Journal of Neurophysiology, 122(4), 1312–1329. 10.1152/jn.00868.2018

6. Bednar, A., & Lalor, E. C. (2020). Where is the cocktail party? Decoding locations of attended and unattended moving sound sources using EEG. NeuroImage, 205, 116283. 10.1016/j.neuroimage.2019.116283

7. Borjigin, A. (2023). The Role of Temporal Fine Structure in Everyday Hearing [Thesis, Purdue University Graduate School]. 10.25394/PGS.19673883.v1

8. Brainard, D. H. (1997). The Psychophysics Toolbox. Spatial Vision, 10(4), 433–436. 10.1163/156856897X00357

9. Broderick, M. P., Anderson, A. J., Di Liberto, G. M., Crosse, M. J., & Lalor, E. C. (2018). Electrophysiological Correlates of Semantic Dissimilarity Reflect the Comprehension of Natural, Narrative Speech. Current Biology: CB, 28(5), 803–809.e3. 10.1016/j.cub.2018.01.080

10. Budescu, D. V. (1993). Dominance analysis: A new approach to the problem of relative importance of predictors in multiple regression. Psychological Bulletin, 114(3), 542–551. 10.1037/0033-2909.114.3.542

11. Campbell, T., Kerlin, J. R., Bishop, C. W., & Miller, L. M. (2012). Methods to Eliminate Stimulus Transduction Artifact From Insert Earphones During Electroencephalography. Ear and Hearing, 33(1), 144. 10.1097/AUD.0b013e3182280353

12. Cooper, J. C., & Gates, G. A. (1991). Hearing in the Elderly-The Framingham Cohort, 1983-1985: Part II. Prevalence of Central Auditory Processing Disorders. Ear and hearing, 12(5), 304–311. 10.1037/h0043158

13. Cowan, N. (2001). The magical number 4 in short-term memory: A reconsideration of mental storage capacity. Behavioral and Brain Sciences, 24(1), 87–114. 10.1017/S0140525X01003922

14. Cowan, N. (2015). George Miller’s Magical Number of Immediate Memory in Retrospect: Observations on the Faltering Progression of Science. Psychological Review, 122(3), 536–541. 10.1037/a0039035

15. Daneman, M., & Carpenter, P. A. (1980). Individual differences in working memory and reading. Journal of Verbal Learning and Verbal Behavior, 19(4), 450–466. 10.1016/S0022-5371(80)90312-6

16. Delorme, A., & Makeig, S. (2004). EEGLAB: An open source toolbox for analysis of single-trial EEG dynamics including independent component analysis. Journal of Neuroscience Methods, 134(1), 9–21. 10.1016/j.jneumeth.2003.10.009

17. Ding, N., & Simon, J. Z. (2012). Emergence of neural encoding of auditory objects while listening to competing speakers. Proceedings of the National Academy of Sciences, 109(29), 11854–11859. 10.1073/pnas.1205381109

18. Dryden, A., Allen, H. A., Henshaw, H., & Heinrich, A. (2017). The Association Between Cognitive Performance and Speech-in-Noise Perception for Adult Listeners: A Systematic Literature Review and Meta-Analysis. Trends in Hearing, 21, 2331216517744675. 10.1177/2331216517744675

19. Füllgrabe, C., & Moore, B. C. J. (2018). The Association Between the Processing of Binaural Temporal-Fine-Structure Information and Audiometric Threshold and Age: A Meta-Analysis. Trends in Hearing, 22, 2331216518797259. 10.1177/2331216518797259

20. Füllgrabe, C., Moore, B. C. J., & Stone, M. A. (2015). Age-group differences in speech identification despite matched audiometrically normal hearing: Contributions from auditory temporal processing and cognition. Frontiers in Aging Neuroscience, 6. https://www.frontiersin.org/articles/10.3389/fnagi.2014.00347

21. Gaudino, E. A., Geisler, M. W., & Squires, N. K. (1995). Construct validity in the trail making test: What makes part B harder? Journal of Clinical and Experimental Neuropsychology, 17(4), 529–535. 10.1080/01688639508405143

22. Getz, L. M., & Toscano, J. C. (2021). The time-course of speech perception revealed by temporally-sensitive neural measures. WIREs Cognitive Science, 12(2), e1541. 10.1002/wcs.1541

23. Gordon-Salant, S., & Fitzgibbons, P. J. (1997). Selected cognitive factors and speech recognition performance among young and elderly listeners. Journal of Speech, Language, and Hearing Research: JSLHR, 40(2), 423–431. 10.1044/jslhr.4002.423

24. Griffiths, T. D., & Warren, J. D. (2004). What is an auditory object? Nature Reviews Neuroscience, 5(11), 887–892. 10.1038/nrn1538

25. Gutschalk, A., Oxenham, A. J., Micheyl, C., Wilson, E. C., & Melcher, J. R. (2007). Human cortical activity during streaming without spectral cues suggests a general neural substrate for auditory stream segregation. The Journal of Neuroscience: The Official Journal of the Society for Neuroscience, 27(48), 13074–13081. 10.1523/JNEUROSCI.2299-07.2007

26. Haier, R. J., Siegel, B. V., Nuechterlein, K. H., Hazlett, E., Wu, J. C., Paek, J., Browning, H. L., & Buchsbaum, M. S. (1988). Cortical glucose metabolic rate correlates of abstract reasoning and attention studied with positron emission tomography. Intelligence, 12(2), 199–217. 10.1016/0160-2896(88)90016-5

27. Hillyard, S. A., Hink, R. F., Schwent, V. L., & Picton, T. W. (1973). Electrical Signs of Selective Attention in the Human Brain. Science, 182(4108), 177–180. 10.1126/science.182.4108.177

28. Hind, S. E., Haines-Bazrafshan, R., Benton, C. L., Brassington, W., Towle, B., & Moore, D. R. (2011). Prevalence of clinical referrals having hearing thresholds within normal limits. International Journal of Audiology, 50(10), 708–716. 10.3109/14992027.2011.582049

29. Holder, J. T., Levin, L. M., & Gifford, R. H. (2018). Speech recognition in noise for adults with normal hearing: Age-normative performance for AzBio, BKB-SIN, and QuickSIN. Otology & Neurotology: Official Publication of the American Otological Society, American Neurotology Society [and] European Academy of Otology and Neurotology, 39(10), e972–e978. 10.1097/MAO.0000000000002003

30. Hopkins, K., & Moore, B. C. J. (2010). Development of a fast method for measuring sensitivity to temporal fine structure information at low frequencies. International Journal of Audiology, 49(12), 940–946. 10.3109/14992027.2010.512613

31. Janse, E. (2012). A non-auditory measure of interference predicts distraction by competing speech in older adults. Aging, Neuropsychology, and Cognition, 19(6), 741–758. 10.1080/13825585.2011.652590

32. Kappenman, E. S., & Luck, S. J. (2011). ERP Components: The Ups and Downs of Brainwave Recordings. In E. S. Kappenman & S. J. Luck (Eds.), The Oxford Handbook of Event-Related Potential Components (p. 0). Oxford University Press. 10.1093/oxfordhb/9780195374148.013.0014

33. Kawahara, H., Masuda-Katsuse, I., & de Cheveigné, A. (1999). Restructuring speech representations using a pitch-adaptive time–frequency smoothing and an instantaneous-frequency-based F0 extraction: Possible role of a repetitive structure in sounds1. Speech Communication, 27(3), 187–207. 10.1016/S0167-6393(98)00085-5

34. Kawahara, H., & Morise, M. (2011). Technical foundations of TANDEM-STRAIGHT, a speech analysis, modification and synthesis framework. Sadhana, 36(5), 713–727. 10.1007/s12046-011-0043-3

35. Kawahara, H., Morise, M., Takahashi, T., Nisimura, R., Irino, T., & Banno, H. (2008). Tandem-STRAIGHT: A temporally stable power spectral representation for periodic signals and applications to interference-free spectrum, F0, and aperiodicity estimation. 2008 IEEE International Conference on Acoustics, Speech and Signal Processing, 3933–3936. 10.1109/ICASSP.2008.4518514

36. Kerlin, J. R., Shahin, A. J., & Miller, L. M. (2010). Attentional Gain Control of Ongoing Cortical Speech Representations in a “Cocktail Party.” Journal of Neuroscience, 30(2), 620–628. 10.1523/JNEUROSCI.3631-09.2010

37. Klein, M., Ponds, R. W. H. M., Houx, P. J., & Jolles, J. (1997). Effect of test duration on age-related differences in stroop interference. Journal of Clinical and Experimental Neuropsychology, 19(1), 77–82. 10.1080/01688639708403838

38. Lu, Z., Daneman, M., & Schneider, B. A. (2016). Does increasing the intelligibility of a competing sound source interfere more with speech comprehension in older adults than it does in younger adults? Attention, Perception, & Psychophysics, 78(8), 2655–2677. 10.3758/s13414-016-1193-5

39. Luck, S. J. (1995). Multiple mechanisms of visual-spatial attention: Recent evidence from human electrophysiology. Behavioural Brain Research, 71(1), 113–123. 10.1016/0166-4328(95)00041-0

40. Luck, S. J. (2022). Applied Event-Related Potential Data Analysis. LibreTexts. 10.18115/D5QG92

41. Lunner, T. (2003). Cognitive function in relation to hearing aid use. International Journal of Audiology, 42 Suppl 1, S49–58. 10.3109/14992020309074624

42. Marcoulides, K. M., & Raykov, T. (2019). Evaluation of Variance Inflation Factors in Regression Models Using Latent Variable Modeling Methods. Educational and Psychological Measurement, 79(5), 874–882. 10.1177/0013164418817803

43. Martin, R. C. (2021). The Critical Role of Semantic Working Memory in Language Comprehension and Production. Current Directions in Psychological Science, 30(4), 283–291. 10.1177/0963721421995178

44. Miller, G. A. (1956). The magical number seven, plus or minus two: Some limits on our capacity for processing information. Psychological Review, 63(2), 81–97. 10.1037/h0043158

45. Miller, L. M., Moore, B. D., & Bishop, C. W. (2020). Frequency-multiplexed speech-sound stimuli for hierarchical neural characterization of speech processing (United States Patent PCT/US15/40629). https://patents.google.com/patent/WO2016011189A1/en

46. Mizumoto, A. (2023). Calculating the Relative Importance of Multiple Regression Predictor Variables Using Dominance Analysis and Random Forests. Language Learning, 73(1), 161–196. 10.1111/lang.12518

47. Moore, B. C. J. (2008). The Role of Temporal Fine Structure Processing in Pitch Perception, Masking, and Speech Perception for Normal-Hearing and Hearing-Impaired People. JARO: Journal of the Association for Research in Otolaryngology, 9(4), 399–406. 10.1007/s10162-008-0143-x

48. Moore, B. C. J. (2021). Effects of hearing loss and age on the binaural processing of temporal envelope and temporal fine structure information. Hearing Research, 402, 107991. 10.1016/j.heares.2020.107991

49. Mueller, S. T., & Piper, B. J. (2014). The Psychology Experiment Building Language (PEBL) and PEBL Test Battery. Journal of Neuroscience Methods, 222, 250–259. 10.1016/j.jneumeth.2013.10.024

50. Nasreddine, Z. S., Phillips, N. A., Bédirian, V., Charbonneau, S., Whitehead, V., Collin, I., Cummings, J. L., & Chertkow, H. (2005). The Montreal Cognitive Assessment, MoCA: A Brief Screening Tool For Mild Cognitive Impairment. Journal of the American Geriatrics Society, 53(4), 695–699. 10.1111/j.1532-5415.2005.53221.x

51. Oberfeld, D., & Klöckner-Nowotny, F. (2016). Individual differences in selective attention predict speech identification at a cocktail party. eLife, 5, e16747. 10.7554/eLife.16747

52. Oh, Y., Bridges, S. E., Schoenfeld, H., Layne, A. O., & Eddins, D. (2021). Interaction between voice-gender difference and spatial separation in release from masking in multi-talker listening environments. JASA Express Letters, 1(8), 084404. 10.1121/10.0005831

53. Oh, Y., Hartling, C. L., Srinivasan, N. K., Diedesch, A. C., Gallun, F. J., & Reiss, L. A. J. (2022). Factors underlying masking release by voice-gender differences and spatial separation cues in multi-talker listening environments in listeners with and without hearing loss. Frontiers in Neuroscience, 16. 10.3389/fnins.2022.1059639

54. Pelli, D. G. (1997). The VideoToolbox software for visual psychophysics: Transforming numbers into movies. Spatial Vision, 10(4), 437–442. 10.1163/156856897X00366

55. Perrone-Bertolotti, M., Tassin, M., & Meunier, F. (2017). Speech-in-speech perception and executive function involvement. PLOS ONE, 12(7), e0180084. 10.1371/journal.pone.0180084

56. Pichora-Fuller, M. K., Schneider, B. A., & Daneman, M. (1995). How young and old adults listen to and remember speech in noise. The Journal of the Acoustical Society of America, 97(1), 593–608. 10.1121/1.412282

57. Picton, T. W., & Hillyard, S. A. (1974). Human auditory evoked potentials. II: Effects of attention. Electroencephalography and Clinical Neurophysiology, 36, 191–200. 10.1016/0013-4694(74)90156-4

58. Reitan, R. M. (1958). Validity of the Trail Making Test as an Indicator of Organic Brain Damage. Perceptual and Motor Skills, 8(3), 271–276. 10.2466/pms.1958.8.3.271

59. Ronnberg, J., Rudner, M., Foo, C., & Lunner, T. (2008). Cognition counts: A working memory system for ease of language understanding (ELU). Int J Audiol, 47 Suppl 2, S99–105. 10.1080/14992020802301167

60. Rudner, M., Lunner, T., Behrens, T., Thorén, E. S., & Rönnberg, J. (2012). Working Memory Capacity May Influence Perceived Effort during Aided Speech Recognition in Noise. Journal of the American Academy of Audiology, 23(08), 577–589. 10.3766/jaaa.23.7.7

61. Ruggles, D., & Shinn-Cunningham, B. (2011). Spatial Selective Auditory Attention in the Presence of Reverberant Energy: Individual Differences in Normal-Hearing Listeners. Journal of the Association for Research in Otolaryngology, 12(3), 395–405. 10.1007/s10162-010-0254-z

62. Schneider, B. A., Rabaglia, C., Avivi-Reich, M., Krieger, D., Arnott, S. R., & Alain, C. (2022). Age-Related Differences in Early Cortical Representations of Target Speech Masked by Either Steady-State Noise or Competing Speech. Frontiers in Psychology, 13. 10.3389/fpsyg.2022.935475

63. Sęk, A. P., & Moore, B. C. J. (2012). Implementation of two tests for measuring sensitivity to temporal fine structure. International Journal of Audiology, 51(1), 58–63. 10.3109/14992027.2011.605808

64. Sęk, A. P., & Moore, B. C. J. (2021). Guide to PSYCHOACOUSTICS. Adam Mickiewicz University Press.

65. Shinn-Cunningham, B. G. (2008). Object-based auditory and visual attention. Trends in Cognitive Sciences, 12(5), 182–186. 10.1016/j.tics.2008.02.003

66. Snyder, J. S., Alain, C., & Picton, T. W. (2006). Effects of attention on neuroelectric correlates of auditory stream segregation. Journal of Cognitive Neuroscience, 18(1), 1–13. 10.1162/089892906775250021

67. Sommers, M. S., & Danielson, S. M. (1999). Inhibitory processes and spoken word recognition in young and older adults: The interaction of lexical competition and semantic context. Psychology and Aging, 14(3), 458–472. 10.1037//0882-7974.14.3.458

68. Stapells, D. (2009). Cortical Event-Related Potentials to Auditory Stimuli. (pp. 395–430).

69. Strauss, E., Sherman, E. M. S., & Spreen, O. (2006). Stroop Test. In A Compendium of Neuropsychological Tests: Administration, Norms, and Commentary (3rd ed., pp. 477–499). Oxford University Press.

70. Strelcyk, O., Zahorik, P., Shehorn, J., Patro, C., & Derleth, R. P. (2019). Sensitivity to Interaural Phase in Older Hearing-Impaired Listeners Correlates With Nonauditory Trail Making Scores and With a Spatial Auditory Task of Unrelated Peripheral Origin. Trends in Hearing, 23, 2331216519864499. 10.1177/2331216519864499

71. Stroop, J. R. (1935). Studies of interference in serial verbal reactions. Journal of Experimental Psychology, 18(6), 643–662. 10.1037/h0054651

72. Teoh, E. S., Ahmed, F., & Lalor, E. C. (2022). Attention Differentially Affects Acoustic and Phonetic Feature Encoding in a Multispeaker Environment. Journal of Neuroscience, 42(4), 682–691. 10.1523/JNEUROSCI.1455-20.2021

73. Teoh, E. S., & Lalor, E. C. (2019). EEG decoding of the target speaker in a cocktail party scenario: Considerations regarding dynamic switching of talker location. Journal of Neural Engineering, 16(3), 036017. 10.1088/1741-2552/ab0cf1

74. The MathWorks Inc. (2021) MATLAB Version: 9.11.0.1769968 (R2021b), Natick, Massachusetts: The MathWorks Inc. https://www.mathworks.com

75. Tremblay, K. L., Pinto, A., Fischer, M. E., Klein, B. E. K., Klein, R., Levy, S., Tweed, T. S., & Cruickshanks, K. J. (2015). Self-Reported Hearing Difficulties Among Adults With Normal Audiograms: The Beaver Dam Offspring Study. Ear and Hearing, 36(6), e290–e299. 10.1097/AUD.0000000000000195

76. Troyer, A. K., Leach, L., & Strauss, E. (2006). Aging and Response Inhibition: Normative Data for the Victoria Stroop Test. *Aging*, Neuropsychology, and Cognition, 13(1), 20–35. 10.1080/138255890968187

77. Unsworth, N., Redick, T. S., Heitz, R. P., Broadway, J. M., & Engle, R. W. (2009). Complex working memory span tasks and higher-order cognition: A latent-variable analysis of the relationship between processing and storage. Memory, 17(6), 635–654. 10.1080/09658210902998047

78. Winkler, I., Denham, S., & Escera, C. (2013). Auditory Event-related Potentials. In D. Jaeger & R. Jung (Eds.), Encyclopedia of Computational Neuroscience (pp. 1–29). Springer. 10.1007/978-1-4614-7320-6_99-1

79. Zekveld, A. A., Rudner, M., Johnsrude, I. S., & Rönnberg, J. (2013). The effects of working memory capacity and semantic cues on the intelligibility of speech in noise. The Journal of the Acoustical Society of America, 134(3), 2225–2234. 10.1121/1.4817926

